# Telomeric DNA breaks in human induced pluripotent stem cells trigger ATR-mediated arrest and telomerase-independent telomere length maintenance

**DOI:** 10.1101/2023.01.19.524780

**Authors:** Katrina N Estep, John W Tobias, Rafael J Fernandez, Brinley M Beveridge, F Brad Johnson

## Abstract

Although mechanisms of telomere protection are well-defined in differentiated cells, it is poorly understood how stem cells sense and respond to telomere dysfunction. Recent efforts have characterized the DNA damage response (DDR) following progressive telomere erosion in human pluripotent cells, yet the broader impact of telomeric double-strand breaks (DSBs) in these cells is poorly characterized. Here, we report on DNA damage signaling, cell cycle, and transcriptome-level changes in human induced pluripotent stem cells (iPSCs) in response to telomere-internal DSBs. We engineered a novel human iPSC line with a targeted doxycycline-inducible TRF1-FokI fusion protein to acutely induce DSBs at telomeres. Using this model, we demonstrate that TRF1-FokI DSBs activate an ATR-dependent DDR in iPSCs, in contrast to an established ATM-dependent response to telomeric FokI breaks in differentiated cells. ATR activation leads to a potent cell cycle arrest in G2, which we show is p53-independent and can be rescued by treatment with an ATR inhibitor. Telomere lengths are remarkably well-maintained in the face of persistent TRF1-FokI induction. Using CRISPR-Cas9 to cripple the catalytic domain of telomerase, we show that telomerase is largely dispensable for survival and telomere length maintenance following telomeric breaks, which instead appear to be repaired by a mechanism bearing hallmarks of lengthening mediated by homologous recombination, so-called alternative lengthening of telomeres (ALT). Our findings suggest a previously unappreciated role for ALT in telomere maintenance in telomerase-positive iPSCs and reveal distinct iPSC-specific responses to targeted telomeric damage.

## Introduction

Telomeres, the protective structures that shield against activation of a DNA damage response (DDR) at chromosome ends, are made up of tandem TTAGGG repeats that are bound by protective shelterin proteins and lengthened by the enzyme telomerase in order to maintain their function [1–3]. In dividing cells, telomere DNA shortens with each round of replication [4,5], until telomeres lose sufficient TTAGGG repeats to support shelterin binding, and telomeres become dysfunctional. Critical telomere erosion in differentiated cells activates a DNA damage response (DDR) that is largely dependent on the ataxia telangiectasia-mutated (ATM) kinase and the downstream activation of the p53 pathway, which promote apoptosis or senescence to halt cellular proliferation [3,6].

However, despite an extensive body of literature on the responses of differentiated cells and cancer cells to telomere dysfunction, the extent to which pluripotent stem cells differ in their telomere capping mechanisms and cellular responses to telomere deprotection remains poorly understood. This is especially of interest considering that pathologies associated with telomere dysfunction, like those observed in telomere-biology disorders, manifest in high-turnover tissues and are linked to stem cell failure[7–11]. It was recently demonstrated that normal chromosome end-protection mechanisms are fundamentally different in pluripotent stem cells than in their differentiated counterparts. Notably, Trf2, the most critical component of the shelterin complex for proper telomere protection in differentiated cells [12], is completely dispensable in murine embryonic stem cells (mESCs) [13,14]. Furthermore, telomere attrition arising from deletion of *hTERT*, the catalytic component of telomerase, has been shown to activate a uniquely ATR-dependent DDR in human iPSCs that differs from the ATM-dependent response that arises in more differentiated cells with critically short telomeres [15,16]. These observations support the concept of a pluripotent stem cell-specific reaction to telomere dysfunction and necessitate additional studies to determine how such cells sense and respond to other forms of telomeric damage.

Here, we address how human induced pluripotent stem cells (iPSCs) respond to DNA double strand breaks (DSBs) within telomere repeats. We engineered a novel human iPSC line harboring an inducible TRF1-FokI allele to acutely induce DSBs within telomeres. Telomeric DSBs activate an ATR-dependent DDR in iPSCs, in contrast to the predominant role of ATM in the response to TRF1-FokI expression in differentiated cells [17]. Induction of TRF1-FokI causes iPSCs to undergo cell cycle arrest in G2, which we show is independent of the downstream DNA damage mediator p53 and can be rescued by treatment with an inhibitor of the ATR kinase. iPSC telomere lengths are remarkably well-maintained following prolonged induction of TRF1-FokI, which we show is largely independent of telomerase. Lastly, we demonstrate that despite having high telomerase activity, iPSCs maintain telomeres after cleavage by the recombination-based alternative lengthening of telomeres (ALT) pathway. Our findings reveal important insights into fundamental telomere maintenance mechanisms in iPSCs and shed light on a previously underappreciated role for telomerase-independent telomere maintenance in human pluripotent stem cells.

## Materials and Methods

### iPSC culture

WT human male iPSC line PENN123i-SV20 was obtained from the University of Pennsylvania iPSC Core Facility. Distribution of this cell line was supported by U01TR001810 from the NIH. iPSCs were grown in mTeSR1 complete medium (STEMCELL Technologies) supplemented with 1% penicillin-streptomycin and maintained at 37°C in the presence of 5% O2 and 5% CO2. iPSCs were cultured on growth factor reduced Matrigel (Corning) coated plates and passaged in clusters every 4-5 days using StemMACS passaging solution. All cells were routinely screened for mycoplasma contamination using a PCR based assay (Uphoff and Drexler, 2014). For single-cell plating, cells were washed once with 1X DPBS (Gibco) and incubated with pre-warmed Accutase (STEMCELL Technologies) for 3 minutes at 37°C, followed by dissociation by pipetting and centrifugation at 200 g for 4 minutes. Cells were then resuspended in mTeSR1 supplemented with 10 μM Thiazovivin (Cayman Chemical) for plating.

### Generation of dox-inducible TRF1-FokI iPSCs

To generate the TRF1-FokI iPSC line, we utilized an established strategy for dox-inducible transgene expression in iPSCs [18]. The TRF1-FokI sequence was obtained from a lentiviral plasmid gifted by Roger Greenberg. DD-ER-TRF1-FokI (lacking the mCherry sequence present in the original construct) was PCR amplified and Gibson cloned into Addgene plasmid #22074 downstream of the TRE promoter in place of the EGFP reporter. For targeted integration of rtTA and DD-ER-TRF1-FokI into the AAVS1 safe harbor locus, vectors encoding a pair of AAVS1-specific zinc finger nucleases (AAVS1-ZFN-L and AAVS1-ZFN-R) and an AAVS1 Neo-CAG-rtTA donor template were obtained from Addgene (plasmids #60915, #60916, and #60431). For transfection, cells were treated with Accutase and seeded at a density of 3.5×10^5 cells per well of a Matrigel-coated 6-well plate to achieve roughly 50% confluency the following day. ZFN-L, ZFN-R, Neo-CAG-rtTA, and sequence-verified Puro-DD-ER-TRE-TRF1-FokI plasmids were transfected into iPSCs using Lipofectamine Stem reagent according to the manufacturer’s protocol. Transfected cells were allowed to recover for 48 hours before being replated as single cells in Matrigel-coated 10cm plates for clonal selection. Cells underwent dual drug selection with 0.25 μg/ml puromycin for 48 hours followed by 40 μg/ml G418 for 10-12 days to select for double targeted clones. Individual clones were expanded, genotyped by PCR using primers targeting both the 5’ and 3’ transgene integration sites, and validated by Sanger sequencing and western blot.

### TRF1-FokI induction

For all experiments unless otherwise stated, TRF1-FokI iPSCs were pre-treated with 1 μg/ml doxycycline for at least 48 hours to allow for maximum expression of TRF1-FokI mRNA. Cells were then induced for the indicated amount of time with 1 μM Shield-1 ligand (Takara bio) to stabilize the TRF1-FokI protein and 1 μM 4-OHT (Cayman Chemical) to promote its nuclear internalization. For all time course experiments including RNA-sequencing, cells were plated at the same density in 6-well plates, pre-treated with dox for 48 hours, induced with Shield-1 and 4-OHT in reverse order (i.e. the longest time point induced first) and collected together at the experiment endpoint for downstream assays.

### Generation of *TP53* KO TRF1-FokI iPSCs

To generate *TP53* knockout TRF1-FokI iPSCs, the gRNA sequence 5’-AATGAGGCCTTGGAACTCA-3 was cloned into the Px330 vector (Addgene plasmid #42230) and transfected into cells using Lipofectamine Stem according to the manufacturer’s protocol. Cells were allowed to recover for 48 hours after transfection before being replated as single cells in a 10 cm dish for clonal selection. Colonies were selected with 5 μM Nutlin-3 for 48 hours. Individual Nutlin-3-resistant colonies were expanded and p53 knockout was confirmed by western blot.

### Generation of biallelic *hTERT* mutant TRF1-FokI iPSCs

To target *hTERT*, 4 different gRNAs were designed to target exon 10 of the *hTERT* catalytic reverse transcriptase domain, cloned individually into the pCAG-SpCas9-GFP-U6-gRNA plasmid (Addgene plasmid #79144), and transfected into cells using Lipofectamine Stem according to the manufacturer’s protocol. Cells were allowed to recover for 40 hours after transfection, then FACS sorted for GFP+ cells and replated in 6-well plates. Individual surviving clones were picked, expanded, and screened for loss of telomerase activity by TRAP assay. More than 50 clones were manually screened, and none showed complete loss of telomerase activity, indicating a strong negative selection against null alleles. The gRNA sequence used to target the sgTERT biallelic mutant clone was 5’TAGGCTGCTCCTGCGTTTGG3’, and successful targeting was confirmed by Sanger sequencing.

### EdU staining using Click-iT™ EdU Imaging Kit

EdU staining was performed using the Invitrogen Click-iT™ EdU Imaging Kit according to the manufacturer’s instructions. Briefly, TRF1-FokI iPSCs were seeded on Matrigel-coated chamber slides in 1 μg/ml dox 48 hours prior to the start of the experiment. Cells were then induced with 1 μM Shield-1 and 1 μM 4-OHT for increasing amounts of time (up to 72 hours). EdU was added at a final concentration of 10 μM for 30 minutes before slides were fixed in 4% PFA, washed, permeabilized, and incubated with 1X Click-iT reaction cocktail containing AlexaFluor-488 for 30 minutes at room temperature protected from light. Following incubation, slides were washed, counterstained with DAPI, air dried, mounted using Prolong Gold Antifade overnight, and imaged on a Leica DM6000 widefield fluorescent scope.

### Propidium iodide and intracellular phospho-H3 flow cytometry

Cells were washed once with 1X DPBS, treated with pre-warmed Accutase to generate single cells, and collected and centrifuged in mTeSR. Pelleted cells were resuspended in 500 ul of 1X DPBS and fixed in 5 ml ice cold 70% EtOH added slowly while vortexing to minimize cell clumping. Cells were allowed to fix overnight at 4°C. For PI staining, cells were pelleted, washed in 1XDPBS, and resuspended in 200 ul of permeabilization solution (0.1% Triton-X in DBPS) containing 10 μl of eBioscience™ propidium iodide (Invitrogen) and 0.5 μg RNase A. Staining and RNA digestion was carried out for 1 hour at room temperature protected from light.

Cells were filtered and analyzed on an Accuri C6 analyzer. For phospho-H3 staining, cells were permeabilized after fixation in 0.1% Triton-X in DPBS for 10 minutes and blocked in 1X DPBS containing 3% BSA and 0.1% Tween-20 for 30 minutes at room temperature. Cells were then incubated with 200 μl of 1XDPBS + 1% BSA + 0.1% Tween-20 containing 4 μl of Alexa Fluor 488-conjugated phospho-histone H3 (Ser10) antibody (Cell Signaling Technologies, 1:50 dilution) for 90 minutes at room temperature, washed, filtered, and analyzed on an Accuri C6 analyzer. All data were quantified using FlowJo software, with manual gating for cell cycle distribution.

### Annexin V and SA ß-galactosidase flow cytometry

Cells were washed once with 1X DPBS, treated with pre-warmed Accutase to generate single cells, and collected and centrifuged in mTeSR. For Annexin V staining, pelleted cells were washed in cold PBS and then resuspended in 1X binding buffer (10 mM HEPES pH 7.4, 140 mM NaCl, 2.5 mM CaCl_2_) at a concentration of 1×10e6 cells/ml. 100 μl of this suspension (1×10e5) was transferred to a 5 ml culture tube and 5 μl of FITC Annexin V (BD Biosciences) added. Cells were stained at room temperature for 30 minutes protected from light and analyzed on an Accuri C6 analyzer using the FL-1 filter setting within 30 minutes of cell collection. SA ß-galactosidase staining was performed using the CellEvent™ Senescence Green Flow Cytometry Assay Kit (Fisher). Single cells were collected, spun down, washed in 1XDPBS, fixed in 2% PFA for 10 minutes, washed, and resuspended in working solution containing 1X Senescence Green Probe diluted in CellEvent™ binding buffer. Cells were incubated for 1 hour at 37°C in the absence of CO_2_ and protected from light. After incubation the working solution was removed and cells were washed with 1% BSA in PBS and analyzed on a BD Accuri C6 analyzer using a 488 nm laser and FL-1 filter setting. All data were quantified using FlowJo software.

### Immunofluorescence staining and microscopy

Cells were seeded on Matrigel-coated chamber slides (Nest Scientific) for immunofluorescence staining. Slides were washed once with 1XDBPS, fixed in 4% paraformaldehyde for 10 minutes at room temperature, washed, permeabilized in 0.5% Triton-X in DPBS for 5 minutes at room temperature, blocked in 3% BSA in 0.1% Triton-X in DPBS for 30 minutes at room temperature, and incubated overnight at 4°C in primary antibody diluted in 1% BSA and 0.1% Tween-20 (for list of antibodies, see Key Resources table). Following primary antibody incubation, slides were washed 3 times in 1XDPBS and incubated in a fluorophore-conjugated secondary antibody for 1 hour at 37°C. Secondary antibody-only slides (no primary) were included as negative staining controls. Following secondary antibody incubation, slides were washed 3 times in PBST (0.1% Tween-20) with 1 μg/ml DAPI added to the penultimate wash. Slides were air dried and mounted in Prolong Gold antifade reagent (Invitrogen) overnight. Images were acquired using either a Nikon Eclipse E600 or a Leica DM6000 widefield fluorescent microscope. Images were prepared and analyzed using Fiji software. Antibodies are listed in Table S1.

### Western blotting

Lysates were prepared by resuspending fresh or previously frozen cell pellets in RIPA lysis buffer (50 mM Tris-HCl pH 7.4, 150 mM NaCl, 1% Triton-X, 1 % sodium deoxycholate, 0.1-0.2% SDS) supplemented with 1X protease inhibitor cocktail (Roche) and 1X phosphatase inhibitor (Cell Signaling Technologies). Cells were lysed on ice for 30 minutes with intermittent vortexing, then centrifuged at 12,000 RPM for 5 minutes at 4°C to pellet cell debris. Supernatant was collected and protein concentration was quantified by Bradford assay using protein assay dye (Bio-Rad) absorbance at 595 nm. 20 μg of protein per sample was resolved on a NuPage 4-12% Bis-Tris polyacrylamide mini gel (Bio-Rad) and transferred to a nitrocellulose membrane in Towbin transfer buffer containing 20% MeOH at 100V for 1 hour or 25V for 16 hours. Membranes were blocked in TBST (1X TBS with 0.1% Tween-20) containing 5% nonfat milk, incubated with primary antibody diluted in blocking solution overnight at 4°C with nutation, rinsed 3 times for 5-10 minutes in TBST, incubated in secondary antibody diluted in blocking solution for 1 hour at room temperature with nutation, and rinsed 3 times for 5-10 minutes in TBST. Membranes were then exposed to Pierce-ECL Western Blotting Substrate (Thermo) and imaged on a Typhoon-FLA scanner. Antibodies are listed in Table S1.

### Cell viability by Cell Titer Blue™ assay

Cell Titer Blue™ reagent was used to measure relative cell viability. Briefly, cells were seeded at equivalent densities in mTeSR supplemented with 10 μM Thiazovivin and 1 μg/ml dox in multiple wells of a Matrigel-coated 24-well plate and allowed to recover for 48 hours before Thiazovivin was withdrawn. Negative control wells were included which contained Matrigel and medium but no cells. Wells were then drugged in triplicate for up to 72 hours with 1 μg/ml dox, 1 μM Shield-1, and 1 μM 4-OHT to induce expression of TRF1-FokI and any other small molecules were added to the assay at the time of induction. At the end of the induction period Cell Titer Blue reagent was added at a final dilution of 1:5 (125 μl per 500 μl of medium in each well), mixed gently, and incubated at 37°C for 2 hours. Fluorescence was recorded at 560/590 nm using a Tecan Infinite 200 Pro plate reader and the average background fluorescence subtracted from each test sample.

### Telomere length measurement by Terminal Restriction Fragment (TRF) analysis

DNA was isolated from cells using a Gentra Puregene core kit A (Qiagen) and quantified by both Nanodrop and quantitative fluorometry using QuBit 2.0 (Invitrogen). For TRF analysis, 500-1000 ng of DNA was digested with CviAII overnight at 25°C followed by digestion with a mixture of BfaI, MseI, and NdeI for 8 hours at 37°C. Southern blotting was carried out using previously established protocols with some modification [19].

Conventional TRF reactions were resolved on a 0.7% agarose gel at 0.833 V/cm for 16-24 hours. For Southern blot analysis, gels were depurinated and denatured and then transferred to a Hybond XL membrane (Cytiva) overnight by capillary transfer using denaturation buffer. The Hybond membrane was hybridized using a DIG-labeled telomere probe at 42°C overnight. The blot was then washed and exposed using CDP-Star on a LAS-4000 Image Quant imager. TRFs were analyzed using ImageQuant.

### Immunofluorescent staining of metaphase chromosomes (Meta-TIF)

Cells were treated with 0.2 μg/ml colcemid for 90 minutes in to arrest cells in metaphase. Colcemid medium was collected and cells were washed once, then dissociated in pre-warmed Accutase, collected into the same conical and pelleted. Supernatant was removed and cells were gently resuspended by flicking in 0.5 ml of residual medium. Cells were swelled in 10 ml of hypotonic solution (0.2% KCl, 0.2% trisodium citrate) at room temperature for 10 minutes and then spun onto poly-L-lysine coated glass slides at 2000 rpm for 10 minutes using a Shandon Cytospin 3. Slides were then fixed in 4% PFA in PBS for 10 minutes at room temperature, washed in 1XDPBS, permeabilized for 10 minutes at room temperature in KCM buffer (120 mM KCl, 20 mM NaCl, 10 mM Tris-HCl pH 7.5) containing 0.1% Triton-X, and blocked with blocking buffer containing 2% BSA and 100 ug/ml RNase A at 37°C for 15 minutes. Slides were incubated with primary antibody for 1 hour at 37°C, washed three times in 1XPBST, incubated with secondary for 1 hour at room temperature, and washed three times again in 1XPBST with DAPI added to the penultimate wash. Slides were then air dried, mounted in Prolong Gold overnight, and imaged using a Leica DM6000 Widefield fluorescent microscope. Images were prepared and analyzed using Fiji software.

### Telomere CO-FISH

For telomere CO-FISH, iPSCs were incubated with 7.5 μM BrdU and 2.5 μM BrdC for 8 hours prior to induction of TRF1-FokI. TRF1-FokI was then induced for 4 hours or mock induced by addition of DMSO. 0.2 μg/ml colcemid was added for the last 2 hours in to arrest cells in metaphase. Medium was collected and cells were washed once, then dissociated in pre-warmed Accutase, collected into the original medium and centrifuged to pellet. Supernatant was removed and cells were gently resuspended by flicking in 0.5 ml of residual medium.

Cells were swelled in 10 ml of prewarmed 0.075 M KCl hypotonic solution at 37°C for 15 minutes and then prefixed by addition of 100 μl 3:1 MeOH/acetic acid fixative solution and pelleted. Metaphase cells were then fixed by washing 3 times in 10 ml of cold 3:1 MeOH/acetic acid and stored at 4°C until ready to spread. Spreading was accomplished by manually dropping fixed metaphase cells onto poly-L-lysine coated glass slides tilted at a 45-degree angle over a steaming beaker of water and blowing on the slides to spread the cells. Slides were air dried overnight at room temperature. On the day of CO-FISH staining, slides were rehydrated in 1XDPBS for 5 minutes, then treated with 0.5 mg/ml RNase A in PBS for 15 minutes at 37°C, rinsed in 1XDPBS, stained with 0.5 μg/ml Hoechst in 2X SSC buffer for 20 minutes at room temperature, and rinsed once more. Slides were then placed in a shallow plastic tray, covered with a thin layer of 2X SSC, and exposed to 365 nm UV using a Stratalinker 1800 UV irradiator set to 5400 μJ x100 at room temperature.

Following UV exposure, BrdU/BrdC-labeled DNA was digested with 10 U/μl Exonuclease III (Promega) in buffer supplied by the manufacturer for 30 minutes at 37°C. Slides were then washed once in 1XDPBS for 5 minutes, fixed in 4% PFA, rinsed in 1XDPBS again, dehydrated in a 70%, 85%, 100% EtOH series for 2 minutes each, and air dried. Hybridization of the first PNA probe (TelC-Cy3) was then carried out in hybridization solution (70% deionized formamide, 0.25% blocking reagent (Roche), 10 mM Tris-HCl pH 7.2) for 2 hours at room temperature. Following the first hybridization, slides were washed twice in PNA wash buffer (70% deionized formamide, 10 mM Tris-HCl pH 7.2, 0.1% BSA). The second hybridization step was carried out by incubating slides with the second PNA probe (TelG-488) diluted in hybridization solution for another 2 hours at room temperature. Lastly, hybridized slides were washed twice with PNA wash buffer for 15 minutes at room temperature on a shaker, washed 3 times with wash buffer II (0.1M Tris-HCl pH 7.2, 0.15M NaCl, 0.08% Tween-20) in a Coplin jar for 5 minutes at room temperature with 1 μg/ml DAPI added to the penultimate wash, air dried, and mounted in Prolong Gold overnight. Stained metaphases were imaged using a Leica DM6000 Widefield fluorescent microscope. Images were prepared and analyzed using Fiji software.

### RNA sequencing

For RNA sequencing, TRF1-FokI iPSCs and parental unedited iPSCs were seeded in triplicate at a density of 25,000 single cells per well of triplicate Matrigel-coated 6-well plates in 1 μg/ml dox and 2 μM Thiazovivin.

Cells were allowed to recover for 48 hours prior to induction, and Thiazovivin was withdrawn on the morning of the second day. TRF1-FokI expression was then induced by addition of 1 μM Shield-1 and 1 μM 4-OHT for 8, 24, or 48 hours, then cells were collected by lysing directly in the plate with TriZol reagent (Fisher) at the experiment endpoint and flash frozen in liquid nitrogen. Total RNA was precipitated from TriZol (Ambion) and cleaned using the Monarch RNA Cleanup Kit (NEB). RNA integrity was verified by the Penn Genomic Analysis Core and only RNAs with RIN>9 were used. RNA library preparation and sequencing was performed by the Penn Next Generation Sequencing Core Facility according to standard protocols. Briefly, high-throughput library prep was performed using Illumina TruSeq stranded mRNA kit and library quality checked on an Agilent Bioanalyzer. Sequencing was performed on a NovaSeq whole S1 flowcell using a 100-cycle kit. For analysis, Salmon was used to count the data against the transcriptome defined in Gencode v41. On a local workstation, several Bioconductor packages in R were used for subsequent steps. The transcriptome count data was annotated and summarized to the gene level with tximeta and further annotated with biomaRt. PCA analysis and plots were generated with PCAtools. Normalizations and statistical analyses were done with DESeq2. GSEA pathway analysis was done against the indicated gene sets from the Molecular Signatures Database (MSigDB), using the DESeq2 statistic as a ranking metric. All GSEA analyses were run with either 10,000 or 20,000 permutations and a weighted enrichment statistic.

### Telomerase Activity Measurement by TRAP Assay

Relative telomerase activity was measured by Telomere Repeat Amplification Protocol (TRAP) assay as previously described [20]. In brief, cells were harvested by Accutase, pelleted, and lysed in 1X CHAPS buffer for 30 minutes on ice. Lysates were then centrifuged at 16,000g for 20 minutes to pellet cell debris and protein concentration was measured by Bradford assay. Serial dilutions of lysate were incubated with a telomerase substrate at 30°C for 30 minutes to allow for telomerase to catalyze the addition of telomere repeats to the substrate. The reactions were then PCR amplified, resolved on a 4-20% TBE polyacrylamide gel and visualized by staining with SYBR Green nucleic acid gel stain. Relative telomerase activity was quantified using ImageJ software.

### C-circle Assay

Genomic DNA was isolated by phenol-chloroform precipitation and quantitated using a Qubit 2.0 fluorometer. C-circle reactions were carried out as previously described [21]. Briefly, 40 ng DNA was reconstituted in 10 ul of TE in a PCR tube, then 9.25 ul of a mastermix containing 2.16X Phi29 DNA polymerase buffer, 0.216% tween-20, 8.65 ug/ml recombinant albumin, and 2 mM each of dATP, dGTP, dTTP, and dCTP was aliquoted to each tube and mixed. U2OS DNA was included as a positive control and HEK293T DNA was included as a negative control. Each assay also included no DNA controls and a set of reactions lacking Phi29 polymerase to account for background. To remove linear genomic DNA, DNA samples were digested with a mixture of CviAII, BfaI, MseI, and NdeI for 16 hours at 25°C followed by 8 hours at 37°C, then digested with 12.5 U λ Exonuclease (NEB) and 100 U Exonuclease I (NEB) for 2 hours at 37°C. Exo-digested DNAs were ethanol precipitated, resuspended in TE, and analyzed on a Southern blot for telomere repeats to confirm complete digestion of linear telomeric DNA. Rolling circle amplification for C-circle detection was carried out at 30°C for 8 hours, followed by a 20-minute incubation at 70°C to inactivate the polymerase. Reactions were then dot blotted onto a Hybond-XL membrane under native conditions, UV-crosslinked, and hybridized using a DIG end-labeled C-rich telomere probe at 42°C overnight. The blot was then blocked, incubated with anti-DIG antibody, washed, and exposed using CDP-Star on a LAS-4000 Image Quant imager.

### Statistical Analysis

Unless otherwise noted, all data is representative of at least three independent experiments conducted on different days using cells of different passage derived from the same engineered clone. Statistical analyses were conducted using GraphPad Prism 9.2 software. Quantitative data is expressed as the mean with error bars representing the standard deviation of the mean. Students t-tests (unpaired and two-tailed) or one-way analysis of variance (ANOVA) were used to determine statistical significance for all comparisons where appropriate. Asterisks indicate the degree of significance (* p<0.05, ** p<0.01, *** p<0.001, **** p<0.0001).

## RESULTS

### A novel iPSC model of inducible telomere dysfunction

To study the impact of telomere dysfunction on pluripotent stem cells, we engineered a human iPSC line in which telomeric double strand breaks (DSBs) can be conditionally induced. Using an established strategy for targeted integration of a doxycycline (dox)-inducible transgene expression system [18], we introduced TRE-DD-ER-TRF1-FokI and CAG-rtTA constructs into the AAVS1 safe harbor loci (**Fig. 1A**). The TRF1-FokI construct contains a destabilization domain (DD) degron and a modified estrogen receptor alpha (ER □) to allow for the controlled stabilization (by Shield-1 ligand) and subcellular localization (by 4-OHT) of a TRF1-FokI fusion protein, which localizes the FokI endonuclease to telomere repeats upon entry into the nucleus [22,23] (**Fig. S1A**). We confirmed that our inducible TRF1-FokI protein was expressed upon addition of dox and Shield-1 ligand (**Fig. 1B**). Immunofluorescence microscopy confirmed a lack of detectable protein expression in the absence of dox and internalization of TRF1-FokI into the nucleus upon addition of 4-OHT (**Fig. 1C**). The TRF1-FokI iPSC line exhibited a normal karyotype (**Fig. S1C**) and retained expression of the transcription factor Nanog, a marker of pluripotency (**Fig. S1E**).

**Fig. 1.**
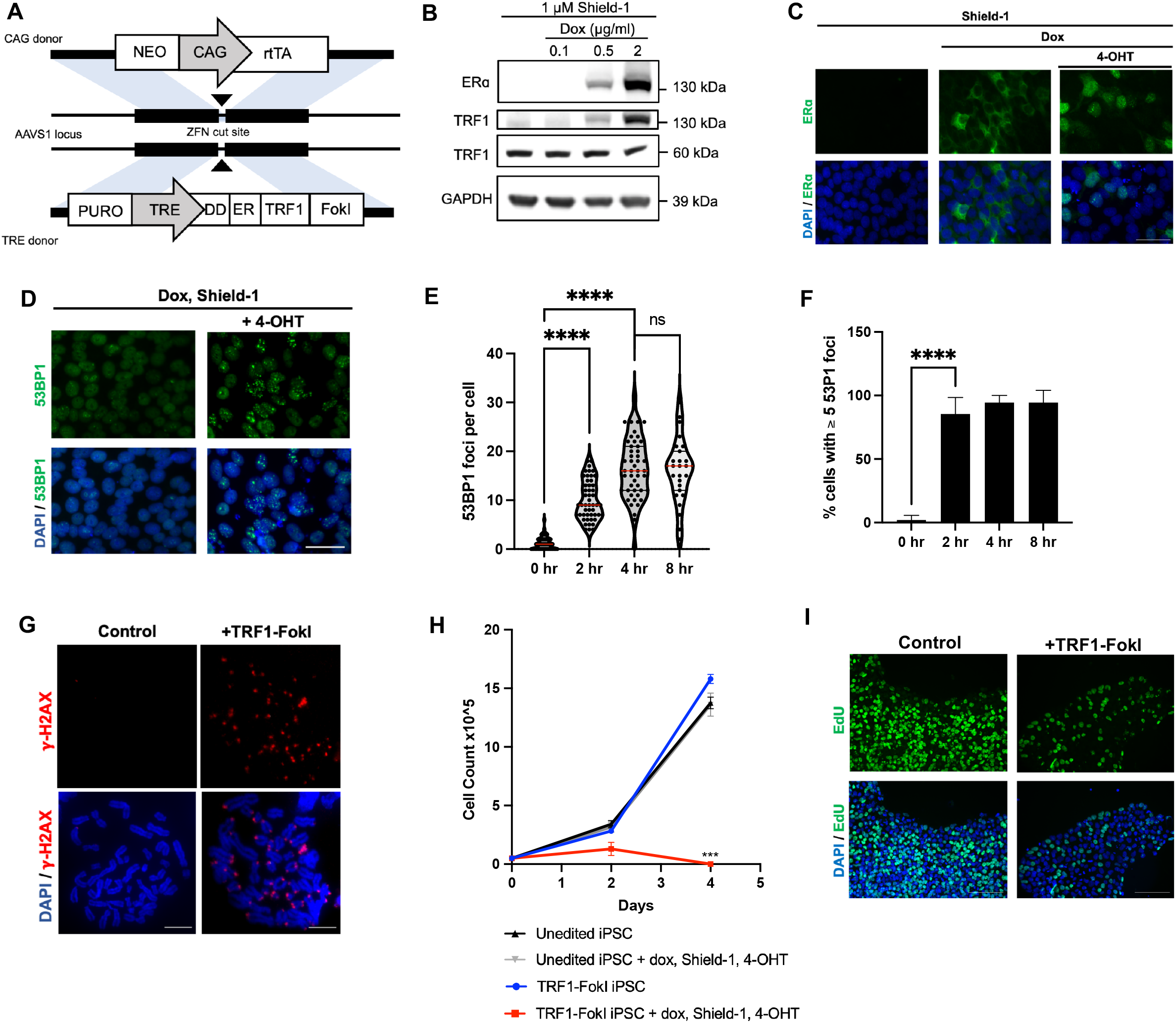
Generation of targeted dox-inducible TRF1-FokI iPSC model. **A)** Strategy for targeted integration of CAG-rtTA and DD-ER-TRF1-FokI into AAVS1 safe harbor loci. **B)** Western blot confirming expression and stabilization of TRF1-FokI fusion protein (130 kDa) using antibodies against ER □ and TRF1. Cells were induced for 48 hours with the indicated concentration of dox and Shield-1. The level of native TRF1 (60 kDa) was unaffected. **C)** Immunofluorescence staining confirming cytoplasmic accumulation of TRF1-FokI fusion protein after 48-hour treatment with dox and Shield-1, and internalization into the nucleus upon addition of 4-OHT for 16 hours. Scale bar=50 μm. **D)** Immunofluorescence staining for 53BP1 foci 2 hours after induction of TRF1-FokI. Scale bar=50 μm. **E)** Quantification of 53BP1 foci as a function of time after nuclear induction of TRF1-FokI. Cells were pre-treated with dox for 48 hours and induced by addition of Shield-1 and 4-OHT. 50-100 nuclei were analyzed per time point across 3 independent experiments. P-values are from one-way ANOVA (p<0.0001). **F)** Quantification of 53BP1 foci as in E but represented as a fraction of all cells exhibiting at least 5 foci. P-value is from one-way ANOVA (p<0.0001). **G)** Immunofluorescence staining for γ-H2AX in metaphase chromosomes harvested from uninduced control cells and cells induced to express nuclear TRF1-FokI for 4 hours. Scale bar=10 μm. **H)** Growth curve for unedited parental and TRF1-FokI edited iPSCs under normal growth conditions and following addition of 1 μg/ml dox, 1 μM Shield-1, and 1 μM 4-OHT. Data are representative of 3 independent experiments. P-value for day 4 is from unpaired two-tailed t-test (p=0.0003). **I)** EdU staining of TRF1-FokI iPSCs under uninduced conditions (control, dox only) and following 72 hours of induction with 1 μM Shield-1 and 1 μM 4-OHT. Scale bar = 100 μm. EdU was added to cultures for 30 minutes prior to fixation.

Previous immunofluorescence microscopy experiments in differentiated cells demonstrated that TRF1-FokI and 53BP1 colocalize at telomeres, indicating a DNA damage response to telomeric breaks [17,22,23]. In our TRF1-FokI iPSCs, we observed a time-dependent accumulation of 53BP1 foci in iPSCs following nuclear induction of TRF1-FokI (**Fig. 1D**), with a significant increase in the average number of 53BP1 foci per cell by 2 hours and a peak at 4 hours post-induction, at which point nearly all cells exhibited more than 5 foci (**Fig. 1E, F**). Prior studies left open the possibility that TRF1-FokI might cause breaks at non-telomeric sites. To verify that the damage was restricted to telomeres, we arrested cells in metaphase and performed immunofluorescent staining for the DNA damage marker γ-H2AX. As expected, cells induced with nuclear TRF1-FokI for 4 hours exhibited metaphase chromosomes with γ-H2AX foci present at chromosome ends (metaphase telomere dysfunction-induced foci, or meta-TIFs, **Fig. 1G, Fig. S2**). Non-telomeric γ-H2AX foci occurred with comparable frequency under uninduced and induced conditions (**Fig. S2D**), confirming that TRF1-FokI selectively cleaves telomeric DNA and does not cut promiscuously throughout the genome.

To gauge the kinetics of iPSC responses to TRF1-FokI, we monitored cell counts and DNA synthesis as indicated by EdU incorporation. TRF1-FokI led to a severe growth defect by 2 days and near complete cell losses by 4 days, as well as diminished EdU incorporation at 72 hours (**Fig. 1H, I**). The lack of any cells surviving beyond 96 hours indicates that most or all cells accumulated toxic levels of TRF1-FokI. Thus, our model allows us to swiftly express TRF1-FokI to selectively induce robust telomeric damage in iPSCs.

### TRF1-FokI DSBs arrest iPSCs in G2 through an ATR-dependent mechanism

Previous work has indicated that human pluripotent cells differ from their isogenic differentiated counterparts in their responses to progressive telomere erosion caused by genetic inactivation of *hTERT* [15]. In particular, iPSCs experiencing critical telomere shortening activated a DNA damage response that is largely dependent upon the ataxia telangiectasia-related (ATR) kinase, rather than ataxia telangiectasia-mutated (ATM) kinase, which is a more prominent mediator of dysfunctional telomere signaling in differentiated cells [17]. We were thus curious which pathway governs responses to acute telomeric DSBs in iPSCs. Western blot analysis across 72 hours of TRF1-FokI induction showed that it led to a notable increase in phosphorylated ATR, but not phosphorylated ATM, suggesting that ATR is a key DNA damage signaling kinase activated in response to telomeric DSBs in iPSCs (**Fig. 2A, B**). Concomitant with ATR activation, we observed an increase in the levels of both Chk1 and Chk2 kinases by 4 hours post-induction of TRF1-FokI (**Fig. 2C**), confirming the activation of a downstream DDR. Flow cytometry for DNA content to evaluate cell cycle dynamics revealed a time-dependent G2/M accumulation in TRF1-FokI iPSCs, with ∼50% of cells arrested in G2/M after 4 hours of induction and >70% by 8 hours (**Fig. 2D, Fig. S3A, B**). Treatment with ATR inhibitor VE821 but not ATM inhibitor KU-55933 at the time of TRF1-FokI induction resulted in a complete rescue of G2/M arrest at 8 hours of induction, consistent with our western analysis indicating a key role for ATR. Note that parental cells (i.e. lacking the TRF1-FokI transgene) treated with induction medium exhibited no change in cell cycle distribution (**Fig. S3C**), indicating the arrest depended on TRF1-FokI.

**Fig. 2.**
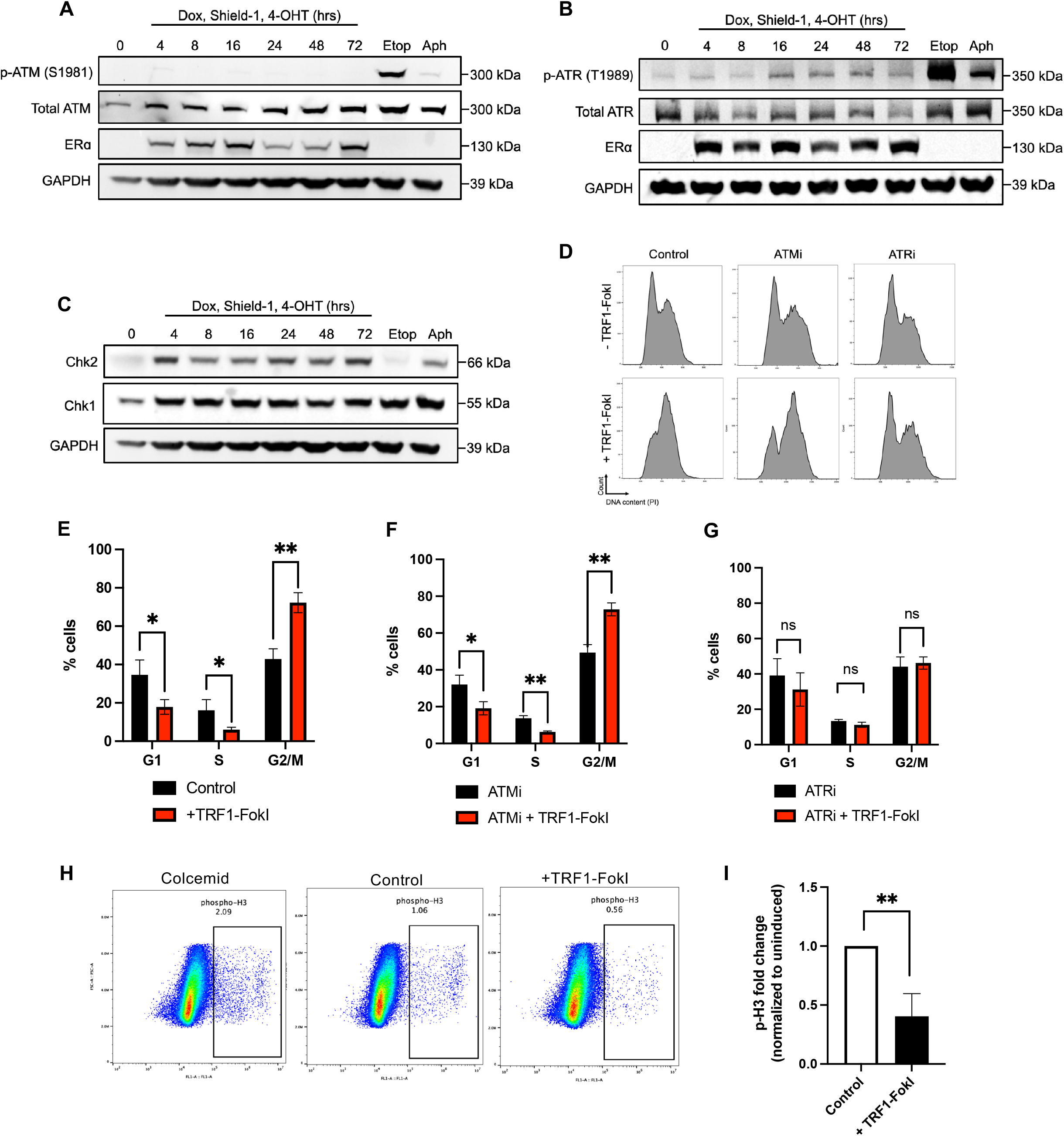
TRF1-FokI DSBs activate ATR signaling to arrest iPSCs in G2. **A, B)** Western blot analysis of ATM and ATR signaling following induction of TRF1-FokI for the indicated amount of time or after treatment with high dose etoposide (10 μM) or aphidicolin (10 μM) for 2 hours. **C)** Effects of ATMi and ATRi on cell cycle following 8 hours of TRF1-FokI induction. Cells were co-treated at the time of TRF1-FokI induction with small molecule ATM inhibitor KU-55933 or ATR inhibitor VE821. DNA content was measured by flow cytometry for propidium iodide. **D-F)** Quantification of cell cycle analyses from three independent experiments carried out as in C. P-values are from unpaired two-tailed t-tests. D) p-values: p=0.028, p=0.039, p=0.002. E) p=0.022, p=0.001, p=0.002. F) p=0.364, p=0.095, p=0.616. **G, H)** Phospho-histone H3 levels in control and TRF1-FokI induced cells as analyzed by flow cytometry (G) and quantification from 3 independent experiments (H). Positive control cells were treated with 0.4 μg/ml colcemid for 1 hour to arrest cells in mitosis. P-value is from unpaired two-tailed t-test (p=0.006).

Telomere erosion in iPSCs was previously shown to trigger mitotic instability, resulting in cell death during mitosis [15]. To examine whether iPSCs experiencing experimental telomeric breaks similarly undergo mitotic arrest, we performed intracellular flow cytometry for phosphorylated histone H3 in TRF1-FokI iPSCs. Surprisingly, at 8 hours of induction, induced cells exhibited on average a two-fold decrease in the percentage of phospho-H3-positive cells compared to control cells, inconsistent with a mitotic arrest phenotype (**Fig. 2H, I**). These data collectively led us to conclude that TRF1-FokI-induced telomeric DSBs arrest iPSCs in G2 by activating an ATR-dependent DDR.

### RNA-seq analysis across a time course of TRF1-FokI induction

Given that pluripotent mouse ESCs lacking the shelterin Trf2 activate specialized gene expression programs required for telomere maintenance compared to their isogenic differentiated counterparts [13], we were next interested in broadly assessing global gene expression changes in TRF1-FokI iPSCs through a time course of TRF1-FokI induction. We performed bulk RNA-sequencing of TRF1-FokI iPSCs at 8, 24, and 48 hours of TRF1-FokI induction compared to uninduced control cells (treated for 48 hours with dox only) as well as compared to the parental unedited iPSC line treated with dox, Shield-1, and 4-OHT for 48 hours (**Fig. 3A**). We found that a sizeable proportion of significantly differentially expressed genes at 48 hours of induction compared to the uninduced cells were also significantly differentially expressed in the parental line (**Fig. S4B, C**), indicating they were driven by exposure to 4-OHT and Shield-1 rather than TRF1-FokI expression, *per se*, and indicating that drug-treated parental cells were the more appropriate control group. In agreement with our cell cycle findings, GSEA analysis of RNA-seq data demonstrated robust activation of ATR but not ATM signaling (**Fig. 3H, I**) and G2/M checkpoint-associated genes in cells expressing TRF1-FokI (**Fig. 3J**).

**Fig. 3.**
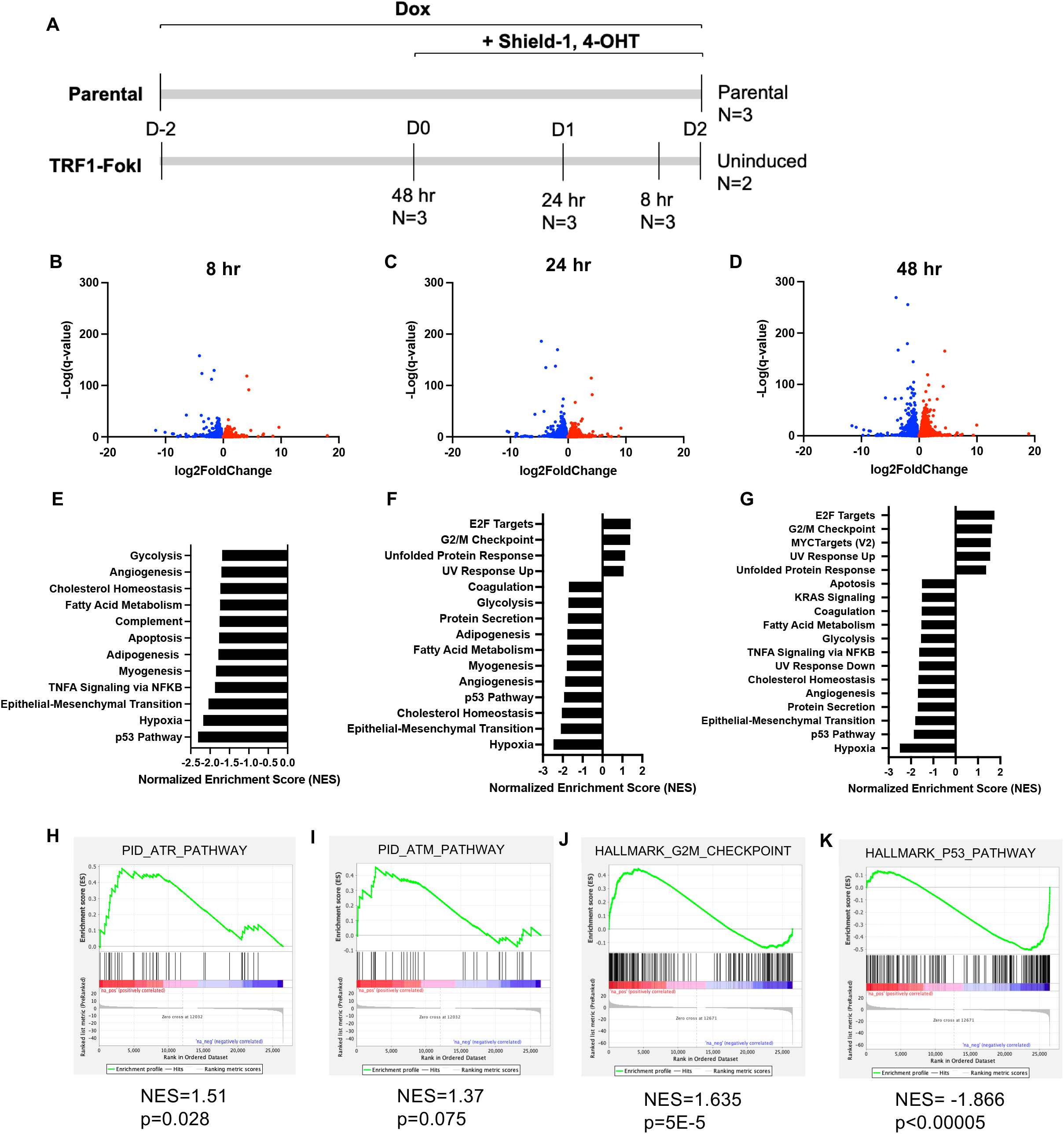
RNA-sequencing reveals time-dependent gene expression changes in TRF1-FokI induced iPSCs. **A)** Experimental setup for RNA-seq experiment. TRF1-FokI cells and unedited parental cells were plated at the same density 48 hours prior to the start of the induction period in medium containing 1 μg/ml dox. Cells were then induced for the indicated amount of time by addition of 1 μg/ml Shield-1 and 1 μg/ml 4-OHT. Multiple replicates per condition were collected together at the experiment endpoint on day 2. Parental cells treated for the duration of the experiment with dox, Shield-1, and 4-OHT serve as a control to account for gene expression changes driven by exposure to Shield-1 and 4-OHT. **B-D**) Volcano plot showing genes significantly (<0.05) upregulated (red) or downregulated (blue) in TRF1-FokI cells induced for 8 (**B**), 24 (**C**), and 48 (**D**) hours compared to unedited parental cells. **E-G**) Gene Set Enrichment Analysis (GSEA) Hallmark gene sets significantly (<0.05) upregulated and downregulated in TRF1-FokI cells induced for 8 (**E**), 24 (**F**), and 48 (**G**) hours compared to unedited parental cells. **H**,**I**) GSEA indicates significant upregulation of ATR pathway (**H**) but not ATM pathway (**I**) at 8 hours of induction compared to unedited parental cells. **J**,**K**) GSEA indicates significant upregulation of G2/M checkpoint (**J**) and significant downregulation of p53 pathway (**K**) gene sets at 48 hours of TRF1-FokI induction compared to unedited parental cells.

### Long-term TRF1-FokI induction leads to non-apoptotic death of iPSCs

Telomere-internal DSBs have been shown to induce differential apoptotic or senescence responses depending on the cell type [24–28]. To assess effects of persistent telomere breaks in iPSCs, we performed a 72-hour induction time course and analyzed cells for markers of viability, apoptosis, and senescence. TRF1-FokI iPSCs showed a time-dependent decline in viability, which was complete between 72 and 96 hours (**Fig. 1H**). The cells remained largely negative for SA ß-gal at all timepoints of induction (<4%, **Fig. S3F**), compared to IMR90 fibroblasts passaged to the point of replicative senescence (>35%). RNA-seq analysis also demonstrated a significant decrease in both p16 and p21 mRNA levels at all three time points of induction (**Fig. S3G**). These observations agree with previous reports that iPSCs do not undergo senescence in response to telomere dysfunction [15]. We also asked whether cells exhibited classic features of apoptosis. There was a progressive increase in annexin V positivity, although this affected only a minority of cells (**Fig. S3E**).

Furthermore, there was no detectable cleaved caspase 3 protein after three days of induction (**Fig. S5A**), suggesting a caspase 3-independent death mechanism, and treatment with the pan-caspase inhibitor Z-VAD-FMK failed to rescue cell growth across the induction period (**Fig. S3I**). Additionally, GSEA hallmarks from RNA-seq showed a significant downregulation of the apoptosis pathway at all three timepoints (**Fig. 3E-G**).

Taken together, these observations suggest that TRF1-FokI cells die by a non-apoptotic mechanism.

### Loss of p53 abrogates cell death without rescuing G2 arrest

To gain a better understanding of DDR signaling in TRF1-FokI iPSCs downstream of ATR, we turned our attention to p53, a key mediator of the DDR. We were surprised to see p53 pathway genes among the most strongly downregulated gene sets at all three time points of TRF1-FokI induction (**Fig. 3E-G, K**). Western blot analysis confirmed that p53 protein levels remained constant throughout 72 hours of TRF1-FokI induction, in contrast to cells treated with high-dose etoposide, in which p53 levels increased approximately twofold within 2 hours (**Fig. S5A**). This observation prompted us to ask whether p53 is required for cell cycle arrest following induction of telomeric DSBs in iPSCs. To test this, we ablated *TP53* using CRISPR-Cas9 in TRF1-FokI iPSCs. The resulting cell line (TRF1-FokI *TP53* KO) exhibited normal karyotype and was resistant to the p53-stabilizing drug Nutlin-3 (**Fig. S5D, E**). Cell cycle analysis revealed that loss of p53 had no effect on G2 accumulation after 8 hours of TRF1-FokI induction (**Fig. 4A-C**), consistent with previous reports that p53 is not required for cell cycle checkpoint activation in iPSCs with telomeres uncapped by progressive erosion [15]. p53 has been reported to regulate cell death through mechanisms distinct from its role in cell cycle checkpoint activation [29]. Consistent with this, we observed that TRF1-FokI *TP53* KO cells were protected against elevated annexin V and cell death after persistent TRF1-FokI induction, with growth kinetics similar to those observed for uninduced WT TRF1-FokI control cells over 72 hours (**Fig. 4D-F**). These results indicate that in iPSCs p53 is not involved in cell cycle checkpoint activation resulting from telomeric DSBs but does regulate cell death following prolonged telomeric damage.

**Fig. 4.**
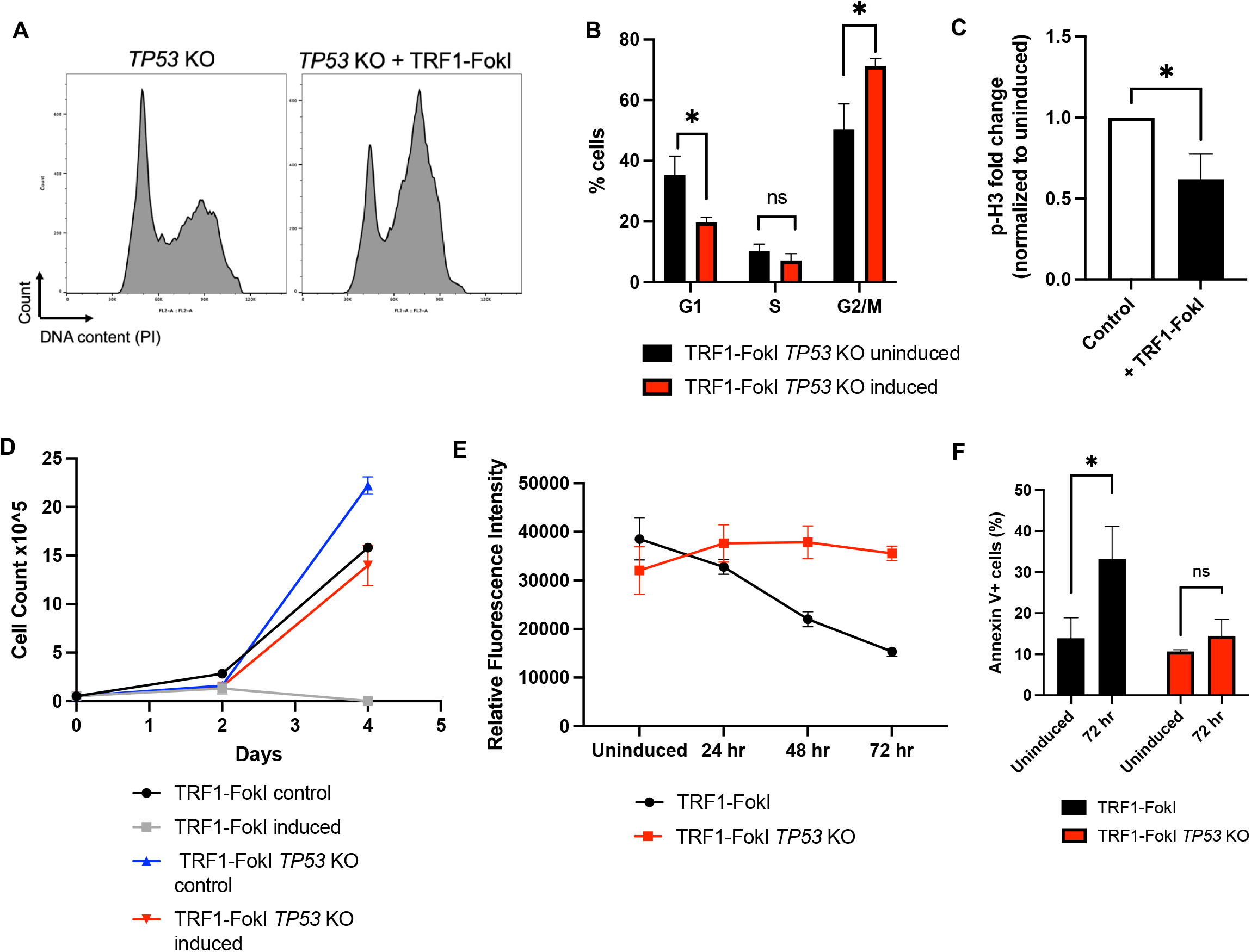
p53 loss attenuates cell death but not G2 arrest following telomeric DSBs in iPSCs. **A)** Cell cycle analysis of control (dox only) and 8-hour induced TRF1-FokI *TP53* KO cells as analyzed by propidium idodide flow cytometry. **B)** Quantification of 3 independent cell cycle analyses carried out as in A. P-values are from unpaired two-tailed t-tests (p=0.014, p=0.179, p=0.015). **C)** Quantification of phospho-H3 levels measured by flow cytometry in p53 KO control cells and *TP53* KO TRF1-FokI induced cells. P-value is from unpaired two-tailed t-test (p=0.013). **D)** Growth curve for *TP53* KO TRF1-FokI iPSCs following induction of TRF1-FokI by addition of 1 μg/ml dox, 1 μM Shield-1, and 1 μM 4-OHT compared to *TP53* WT parental TRF1-FokI engineered line. Data are representative of 3 independent experiments. **E)** Fluorometric cell viability assay measuring total cell viability of parental TRF1-FokI or TRF1-FokI *TP53* KO cells over 72 hours of TRF1-FokI induction. Data represent the mean and SD from triplicate samples measured per time point per cell line. **F)** Quantification of annexin V positivity in TRF1-FokI *TP53* KO cells before induction and after 72 hours of induction as analyzed by flow cytometry and quantified from 3 independent experiments. P-values are from unpaired two-tailed t-tests (p=0.022, p=0.06).

### Human iPSCs do not activate a totipotent-like two-cell-stage transcriptional program in response to telomeric DSBs

mESCs are capable of adopting a 2C-like gene expression state characterized by high expression of the *Zscan4* gene cluster, which is critical for telomere protection in the absence of the shelterin Trf2 [13]. To delineate differences between mouse vs. human pluripotent cells, as well as the nature of responses to acute telomeric DSBs vs. Trf2 deletion, we examined whether 2C-like genes were significantly altered in our system following TRF1-FokI expression. RNA-seq showed that *ZSCAN4* transcripts were not enriched at any of the induced time points (**Fig. S6A, B**), and neither were transcripts mapping to the transcription factor *DUX4* (data not shown), which in mouse drives expression of *Zscan4* and regulates the 2C state in mESCs [30,31].

Additionally, GSEA confirmed that ZSCAN4 transcriptional targets were not enriched at any of the three time points (**Fig. S6C**). We next asked whether human orthologues of murine 2C genes previously shown to be upregulated following *Trf2* deletion in mESCs were similarly upregulated in our system. Of the 13 other 2C genes previously reported, 7 have conserved human orthologues, 6 of which we detected by RNA-seq; of those 6, only 2 were significantly upregulated at 48 hours of induction (**Fig. S6A**). We thus conclude that human iPSCs do not activate a ZSCAN4-dependent 2C-like transcriptional state in response to targeted telomeric DSBs.

### TRF1-FokI breaks result in negligible telomere shortening and undergo telomerase-independent length maintenance

We next asked how persistent expression of TRF1-FokI impacts telomere length maintenance. We hypothesized that prolonged induction of TRF1-FokI would lead to a time-dependent decrease in bulk telomere length as telomeres are cleaved, as has been shown to be the case for MEFs [17]. Surprisingly, persistent expression of TRF1-FokI in iPSCs resulted in negligible telomere shortening as analyzed by terminal restriction fragment (TRF) analysis (**Fig. 5A**). This was unexpected, given that 53BP1 and γ-H2AX staining indicated widespread telomere cleavage (**Fig. 1**). Compared to differentiated cells, iPSCs have long telomeres maintained by high levels of *hTERT* and robust telomerase activity [32]. We therefore hypothesized that the lack of telomere shortening was a result of efficient extension of cleaved telomeres by telomerase with kinetics comparable to the rate of cleavage. We first asked whether we could detect elevated telomerase activity in response to TRF1-FokI breaks, as other DSB-inducing agents (including ionizing radiation and etoposide) have been shown to boost telomerase expression and enzymatic activity in various cellular contexts [33,34]. Our RNA-seq data was consistent with this idea, as the telomerase RNA component (*TERC*) showed significantly increased expression at 24 and 48 hours (**Fig. 5B, C**). However, a functional test for telomerase enzymatic activity by TRAP assay showed no detectable change in TRAP activity with TRF1-FokI induction (**Fig. 5E, F**), indicating that TRF1-FokI breaks do not promote increased telomerase activity. To directly test the contribution of telomerase in maintaining telomere lengths following FokI breaks, we used CRISPR-Cas9 to target the catalytic domain of *hTERT* in TRF1-FokI iPSCs. We obtained a surviving hypomorph clone (sgTERT) with biallelic indel mutations within the reverse transcriptase domain that severely compromised telomerase activity (**Fig. 5G, H, Fig. S7**). sgTERT cells displayed reduced growth kinetics at baseline compared to the parental TRF1-FokI line (**Figure 5I**), but taking this into account, the rate of impaired growth following induction of TRF1-FokI was no more severe for sgTERT cells than it was for cells with WT *TERT* (**Fig. 5J**). Furthermore, although TRF analysis of sgTERT cells revealed obvious telomere shortening during the ∼25 divisions since its generation (**Cf. Fig. 5K vs. 5A**), additional telomere shortening was not apparent over 3 days of TRF1-FokI induction. Collectively, these data led us to conclude that telomere lengths are maintained following persistent TRF1-FokI breaks through a largely telomerase-independent mechanism.

**Fig. 5.**
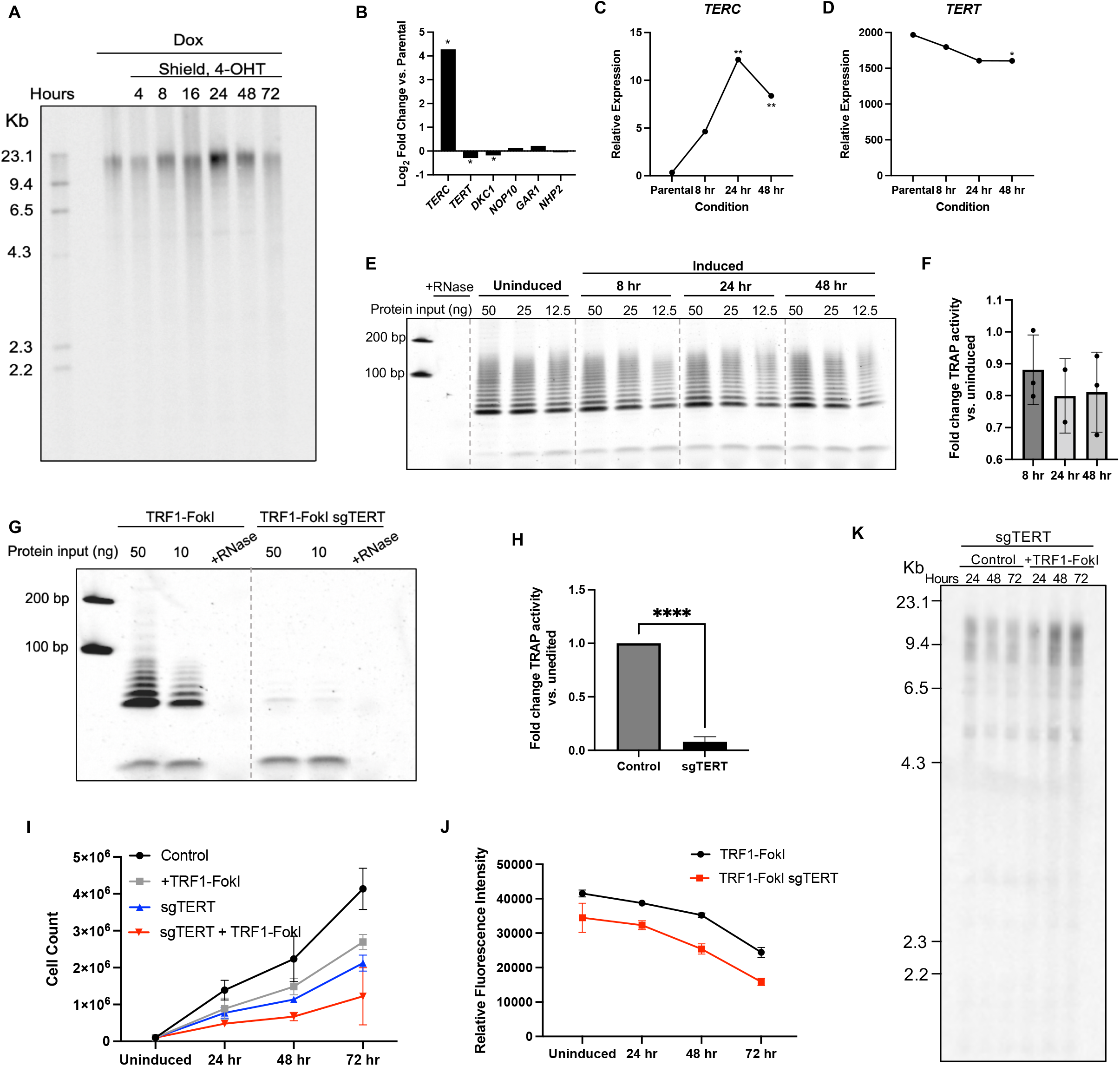
Telomere length is well-maintained following prolonged induction of TRF1-FokI. **A)** Terminal Restriction Fragment (TRF) analysis of telomere lengths in uninduced cells and induced to express TRF1-FokI for the indicated number of hours. **B)** mRNA expression levels of components of the telomerase holoenzyme from RNA-seq at 48 hours of induction compared to uninduced parental cells. *p-value<0.05. **C)** Trend of relative mRNA expression of telomerase RNA component (*TERC*) at each of the indicated time points as measured by RNA-seq. **p-value<0.01. **D)** Trend of relative mRNA expression of telomerase reverse transcriptase (*TERT*) at each of the indicated time points as measured by RNA-seq. *p-value<0.05. **E)** Telomerase repeat amplification protocol (TRAP) assay measuring telomerase enzymatic activity in TRF1-FokI iPSCs using twofold extract dilutions. **F)** Quantification of fold change TRAP activity of 8, 24, and 48-hour induced TRF1-FokI iPSCs compared to uninduced cells. **G)**TRAP assay measuring telomerase enzymatic activity in uninduced TRF1-FokI iPSCs and biallelic *hTERT* mutant TRF1-FokI iPSCs (sgTERT) using fivefold extract dilutions. **H)** Quantification of fold change in TRAP activity of biallelic *hTERT* mutant TRF1-FokI iPSCs (sgTERT) compared to parental TRF1-FokI iPSCs. Data represent mean and SD from two independent experiments. p<0.0001. **I)** Growth curve for TRF1-FokI (WT *hTERT*) and biallelic *hTERT* mutant TRF1-FokI iPSCs (sgTERT). All cells were treated with 1 μg/ml dox for the duration of the experiment, and cells were either mock induced (control, dox only) or induced (1 μg/ml dox, 1 μM Shield-1 and 1 μM 4-OHT) for the indicated duration, then total cells were counted at each time point. **J)** Fluorometric cell viability assay measuring total cell viability of parental TRF1-FokI or biallelic *hTERT* mutant TRF1-FokI (sgTERT) cells over 72 hours of TRF1-FokI induction. Data represent the mean and SD from 6 samples measured across two plates per time point per cell line. **K)** TRF analysis of telomere lengths in biallelic *hTERT* mutant (sgTERT) TRF1-FokI iPSCs either mock induced (control, left) or induced to express TRF1-FokI (+TRF1-FokI) for the indicated number of hours.

Telomere recombination is central to the telomerase-independent ALT pathway. Mouse ESCs can employ ALT [35–37], but whether it can occur following acute telomeric breaks in human iPSCs has not been tested previously. To begin to examine this possibility, we conducted a focused analysis of RNA sequencing data for major DNA repair pathways. Genes associated with homology-directed repair (HDR) were generally upregulated, including *RAD51* and its paralogs *RAD51B* and *RAD51C*, as well as *BLM, POLD3, BRCA1* and *BRCA2*. (**Fig. 6A, C**), and this was corroborated by GSEA analysis (**Fig. 6B**). Importantly, there was no enrichment of genes associated with the microhomology-mediated/alternative non-homologous end joining pathway (MMEJ/alt-NHEJ), including *LIG3* or *PARP1*, which is a major mechanism by which telomeric FokI breaks are repaired in differentiated cells [17] (**Fig. 6B, C**).

**Fig. 6.**
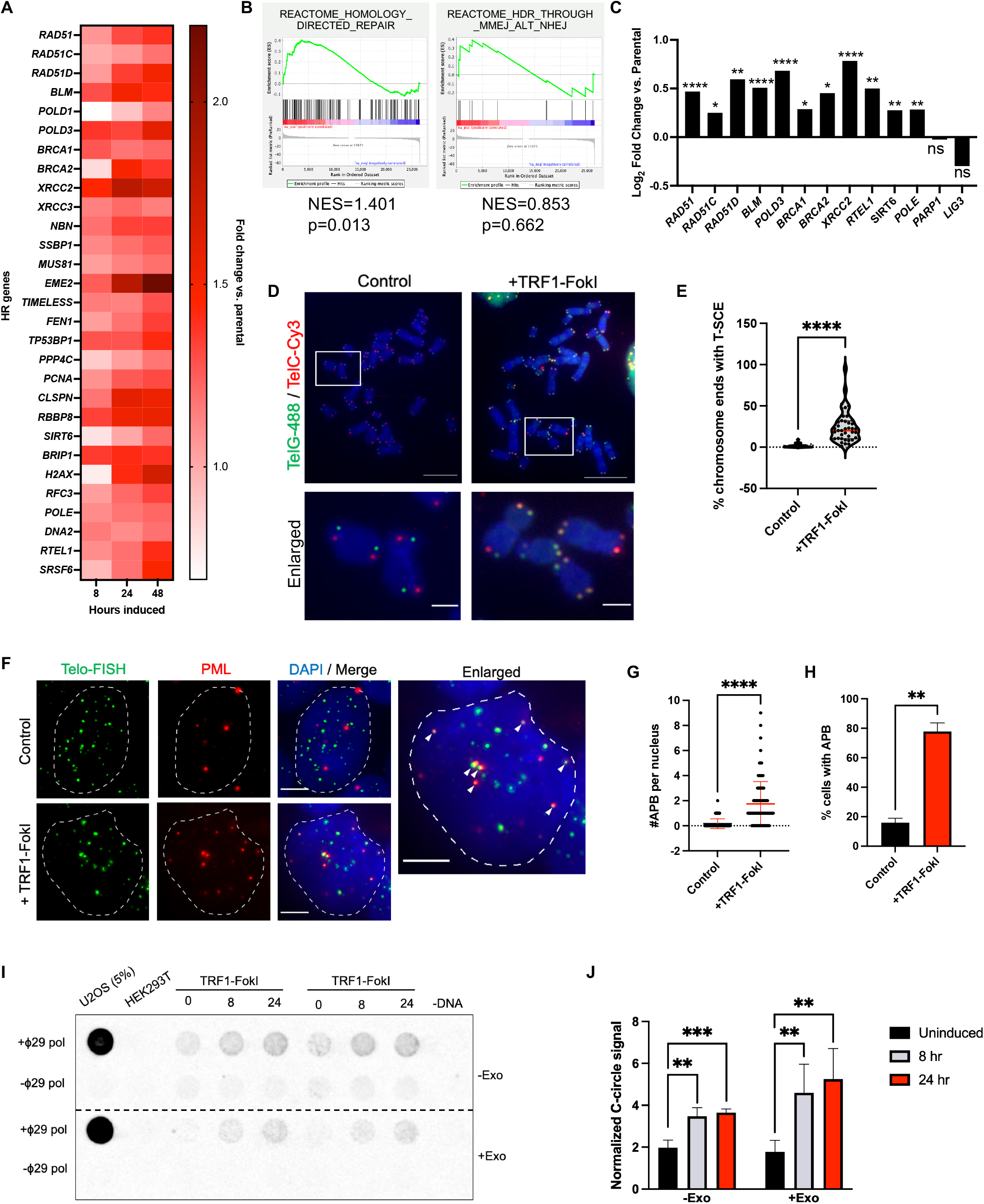
TRF1-FokI DSBs activate ALT in iPSCs. **A)** Heat map depicting fold change in relative expression normalized to unedited parental cells of HR-related genes throughout the 48-hour induction period. **B)** GSEA indicating a significant enrichment of the homology-direct repair reactome gene signature, but not the microhomology-mediated / alternative end-joining signature, at 48 hours of TRF1-FokI induction. **C)** Significant upregulation (measured by Log2(fold change) vs. unedited parental cells) of many genes associated with homologous recombination at 48 hours of TRF1-FokI induction. **D)** Chromosome orientation (CO)-FISH performed on metaphase spreads harvested from uninduced (left, dox only) and 4-hour induced (right) TRF1-FokI iPSCs. Leading strand telomeres were hybridized with a 488-conjugated TelG PNA FISH probe (green) and lagging strand telomeres were hybridized with a Cy3-conjugated TelC PNA FISH probe (red). Top panels: scale bar=10 μm. Enlarged panels: scale bar=2 μm. **E)** Quantification of T-SCE frequency as detected by CO-FISH. Data are representative of 3 independent experiments. **F)** Combined IF-FISH staining for telomeric DNA (green) and PML protein (red) in uninduced control (top panels) and 48-hour induced (bottom panels) TRF1-FokI iPSCs. Arrows indicate representative ALT-associated PML bodies (APBs). Scale bar=5 μm. **G)** Quantification of discrete number of APBs per nucleus in uninduced control and 48-hour induced TRF1-FokI iPSCs. Data are representative of at least 100 nuclei across 3 independent experiments. p<0.0001 from unpaired two-tailed t-test. **H)** Quantification of the fraction of cells counted containing one or more APBs in uninduced control and 48-hour induced TRF1-FokI iPSCs. Data are representative of at least 100 nuclei across 3 independent experiments. p=0.006 from unpaired two-tailed t-test. **I)** Representative dot blot from C-circle assay (CCA) performed using genomic DNA harvested from uninduced, 8-hour induced, and 24-hour induced TRF1-FokI iPSCs (N=2 biological replicates shown). ALT+ U2OS and ALT-HEK293T DNAs were included as positive and negative controls, respectively. CCA was performed on bulk genomic DNA (top) and following removal of linear DNA by exonuclease digestion (bottom) to confirm circularity of template. Sample-specific background is quantified from signal generated in the absence of ϕ29 polymerase. **J)** Quantification of relative C-circle abundance in uninduced (0 hr) and induced (8 hr, 24 hr) TRF1-FokI iPSCs with and without exonuclease treatment to remove linear genomic DNA. C-circle signal for each sample is normalized to sample-specific background. Data represent average and SD from 2 technical replicates and 2 biological replicates (N=4).

To test whether TRF1-FokI cells exhibit *bona fide* features of ALT, we first carried out chromosome orientation (CO)-FISH, which can be used to visually detect telomeric sister chromatid exchange events (T-SCEs) [38,39]. We detected a significant increase in the frequency of T-SCEs in cells induced to express TRF1-FokI for 4 hours (**Fig. 6D, E**), confirming active recombination between telomeres. Immunofluorescent staining also revealed a significant increase in the colocalization of PML protein with TTAGGG repeats (**Fig. 6F, G**), which is characteristic of ALT cells [40,41]. Furthermore, we probed for ALT-specific C-rich telomeric DNA circles (C-circles) by rolling circle amplification [21,42]. C-circles were present in TRF1-FokI iPSCs, and they became more abundant with induction (**Fig. 6I, J**). The abundance of C-circles in TRF1-FokI iPSCs was low compared to ALT+ U2OS cells, raising the question as to whether the signal might be from spurious amplification of nicked linear telomeric DNA rather than C-circles. However, the signal remained after restriction digestion and exonuclease treatment to remove linear genomic DNA, confirming that the signal was amplified entirely from circular template. Taken together, these results indicate that telomeric DSBs activate ALT in telomerase-positive iPSCs.

## DISCUSSION

We engineered a human iPSC line with an inducible TRF1-FokI construct to study the impact of acute telomeric DSBs on iPSC growth, survival, DNA damage signaling, and repair mechanisms. Protein and cell cycle analyses showed that such breaks activate ATR signaling to arrest cells in G2. Importantly, this finding differs from the primarily ATM-dependent response to the same form of targeted telomeric DSBs in differentiated cells [17]. Since ALT takes place exclusively in G2, it is tempting to speculate that ATR rather than ATM activation in this context serves to promote telomere recombination, which could use an intact telomere on one sister chromatid as a template for repair of a broken sister chromatid telomere as a way to achieve efficient and error-free restoration of telomere length. This would be a particularly useful telomeric repair mechanism for pluripotent stem cells, which must preserve genomic integrity in order to faithfully serve as the progenitors of an entire organism.

Our finding that iPSCs do not undergo senescence or canonical apoptosis in response to targeted telomeric DSBs is also in stark contrast to what is known to occur in other cell types. In differentiated cells, prolonged cell cycle checkpoint activation engages downstream apoptosis or senescence to halt proliferation of genetically unstable cells. While additional studies are needed to pinpoint the precise mechanism by which our TRF1-FokI iPSCs die, we hypothesize that a necroptosis-like mechanism might be at play, as it is associated with annexin V positivity but not caspase activation [43]. Necroptosis has been tightly linked to upregulation of the unfolded protein response (UPR) pathway [44–46], which our RNA-seq detected was significantly elevated at later timepoints of TRF1-FokI induction (**Fig. 3**). As an aside, we and others have previously observed upregulation of the UPR in the setting of telomere dysfunction in other cellular contexts [47,48], suggesting that telomere damage may be a general driver of ER stress.

It is noteworthy that we were unable to obtain iPSCs that are fully null for *hTERT*. This differs from the case for murine ESCs and iPSCs, and presumably reflects the well-established greater susceptibility of humans to telomerase deficiency (*e*.*g*., first-generation mice null for telomerase activity are nearly normal [49,50], whereas people with hypomorphic alleles in telomerase components can exhibit severe defects in childhood [51,52]). It also appears to contrast with a recent report that *hTERT* could be deleted fully in human iPSCs [15], but these cells contained a dox-inducible *hTERT* transgene which was expressed during generation of the endogenous *hTERT* knockout and thus presumably maintained viability. An essential role for telomerase in human iPSCs may reflect one or more of the apparent extra-telomeric roles for telomerase [53,54], consistent with the poor growth of our hypomorphic clone even at relatively long telomere lengths. It is also remarkable that telomeres shortened with cell division in our hypomorphic sgTERT clone, but telomere lengths were nonetheless maintained following cleavage with TRF1-FokI. This indicates that HDR can more readily repair telomeres with DSBs within the telomere repeat than the gradually shortened telomere termini that result from the so-called end-replication problem. Importantly, such internal telomere breaks occur naturally, *e*.*g*. during DNA replication of telomeres [55], and so it is not surprising that mechanisms for their repair have evolved that can distinguish them from the gradual shortening caused by the end-replication problem.

Murine Zscan4 is a critical regulator of HDR-based telomere maintenance in mESCs [13,36,56–58]. However, we found no evidence that human iPSCs activate a ZSCAN4-dependent transcriptional state in response to targeted telomeric damage. We speculate that this failure to activate ZSCAN4 or other 2C-like genes may reflect a greater similarity of iPSCs to the “primed” mESCs that are derived from postimplantation epiblast, rather than “naïve” mESCs derived from the inner cell mass, and that acquisition of the 2C state may be restricted to naïve pluripotency, though this remains to be tested. Additionally, several key distinctions between our experimental system and others may contribute to this difference: murine vs. human, ESC vs. iPSC, and progressive telomere shortening vs. acute internal telomere breaks; further studies will be required to better understand the full range of transcriptional responses to telomere dysfunction across different stem cell types.

We found that TRF1-FokI iPSCs undergo negligible telomere shortening over several days of TRF1-FokI induction, irrespective of telomerase enzymatic activity, and activate features of ALT in response to telomeric DSBs. Very little is known regarding the importance of ALT in maintaining iPSC telomere length under normal culture conditions and following damage. It has been shown, for example, that murine ESC and iPSC telomeres undergo transient ALT-dependent lengthening during the derivation and reprogramming processes [35–37,59], but it was previously unclear whether ALT is employed by human iPSCs and whether fully reprogrammed iPSCs, either murine or human, utilize ALT or rely exclusively on telomerase to maintain telomeres. The fact that we observe low levels of APBs and C-circles in our cells even under uninduced conditions suggests that ALT may play a heretofore underappreciated role in the homeostatic regulation of telomere length in pluripotency, and our finding that features of ALT significantly increase following acute telomeric DSBs underscores that recombination is an important component of the repair of iPSC telomeres. Furthermore, our findings argue against the concept of a given cell type being strictly “ALT+” or “ALT-” and instead suggest that ALT may serve as an important telomere maintenance mechanism in cells that are normally characterized by high telomerase activity. We speculate that this is a unique characteristic of pluripotent stem cells that makes them remarkably adept at both maintaining telomere length and repairing telomeric damage, owing to their requirement for adequately long and well-protected telomeres to retain their functionality as stem cells; to be able to proliferate indefinitely and give rise to all cellular lineages. Taken together, our study highlights that iPSCs have specialized responses to telomeric damage and raises important implications for their use as a model to understand normal development, telomere-biology disorders, cancer, tissue regeneration, and cellular aging.

## ACKNOWLEDGEMENTS

We would like to thank Dr. Roger Greenberg for providing the TRF1-FokI construct and Drs. Chris Lengner, Craig Bassing, Kathryn Hamilton, and Elizabeth Furth for fruitful discussions. We would also like to thank Hunter Reavis for insightful editorial suggestions. This work was supported in part by the following core facilities at the University of Pennsylvania: Induced Pluripotent Stem Cell Core, Cell and Developmental Biology Microscopy Core, Molecular Pathology and Imaging Core, Next Generation Sequencing Core, Genomic Analysis Core, and Cell Center Stockroom; as well as by the Human Pluripotent Stem Cell Core and Flow Cytometry Core Laboratory at the Children’s Hospital of Philadelphia.

## DECLARATIONS

### Funding

This work was supported by the NIH (R01HL148821 and F31CA260918) and the Patel Family Cancer Research Scholar Award from the University of Pennsylvania Abramson Cancer Center.

### Disclosure of conflicts of interest

The authors have no financial or non-financial conflicts to disclose.

### Ethics approval

Not applicable.

### Consent to participate

Not applicable.

### Consent for publication

Not applicable.

## Availability of data and material

Data generated and/or analyzed during the current study are available from the corresponding author upon request. RNA-seq data will be deposited in GEO.

## Authors’ contributions

KNE and FBJ conceived the study. KNE conducted all experiments and analyses and wrote and revised the manuscript with editorial suggestions from RJF and FBJ. JWT conducted bioinformatics analyses related to RNA-seq data. BMB assisted with experiments. FBJ supervised the project. All authors read and approved the final manuscript.

## Notes

### Competing Interest Statement

The authors have declared no competing interest.

## References

1. Greider, C. W., & Blackburn, E. H. (1985). Identification of a specific telomere terminal transferase activity in Tetrahymena extracts. Cell, 43(2 Pt 1), 405–413. https://linkinghub.elsevier.com/retrieve/pii/0092867485901709

2. de Lange, T. (2005). Shelterin: the protein complex that shapes and safeguards human telomeres. Genes & Development, 19(18), 2100–2110. http://www.genesdev.org/cgi/doi/10.1101/gad.1346005

3. Shay, J. W., & Wright, W. E. (2019). Telomeres and telomerase: three decades of progress. Nature Reviews. Genetics, 20(5), 299–309. http://www.nature.com/articles/s41576-019-0099-1

4. Olovnikov, A. M. (1971). [Principle of marginotomy in template synthesis of polynucleotides]. Doklady Akademii Nauk SSSR, 201(6), 1496–1499. http://eutils.ncbi.nlm.nih.gov/entrez/eutils/elink.fcgi?dbfrom=pubmed&id=5158754&retmode=ref&cmd=prlinks

5. Biology, JD Watson Nature New, & 1972. (n.d.). Origin of concatemeric T7DNA. Springer. https://link.springer.com/content/pdf/10.1038/newbio239197a0.pdf

6. d’Adda di Fagagna, F., Reaper, P. M., Clay-Farrace, L., Fiegler, H., Carr, P., Von Zglinicki, T., Saretzki, G., Carter, N. P., & Jackson, S. P. (2003). A DNA damage checkpoint response in telomere-initiated senescence. Nature, 426(6963), 194–198. http://www.nature.com/articles/nature02118

7. Armanios, M., Alder, J. K., Parry, E. M., Karim, B., Strong, M. A., & Greider, C. W. (2009). Short telomeres are sufficient to cause the degenerative defects associated with aging. The American Journal of Human Genetics, 85(6), 823–832. https://doi.org/10.1016/j.ajhg.2009.10.028

8. Hao, L.-Y., Armanios, M., Strong, M. A., Karim, B., Feldser, D. M., Huso, D., & Greider, C. W. (2005). Short telomeres, even in the presence of telomerase, limit tissue renewal capacity. Cell, 123(6), 1121–1131. https://doi.org/10.1016/j.cell.2005.11.020

9. Lee, H. W., Blasco, M. A., Gottlieb, G. J., Horner, J. W., 2nd, Greider, C. W., & DePinho, R. A. (1998). Essential role of mouse telomerase in highly proliferative organs. Nature, 392(6676), 569–574. https://doi.org/10.1038/33345

10. Nag, S. (2020). Syndromes associated with telomere shortening. In Telomerase and non-Telomerase Mechanisms of Telomere Maintenance. IntechOpen. https://doi.org/10.5772/intechopen.88792

11. Armanios, M. (2022). The role of telomeres in human disease. Annual Review of Genomics and Human Genetics, 23(1), 363–381. https://doi.org/10.1146/annurev-genom-010422-091101

12. van Steensel, B., Smogorzewska, A., & de Lange, T. (1998). TRF2 protects human telomeres from end-to-end fusions. Cell, 92(3), 401–413. https://linkinghub.elsevier.com/retrieve/pii/S0092867400809320

13. Markiewicz-Potoczny, M., Lobanova, A., Loeb, A. M., Kirak, O., Olbrich, T., Ruiz, S., & Lazzerini Denchi, E. (2021). TRF2-mediated telomere protection is dispensable in pluripotent stem cells. Nature, 589(7840), 110–115. https://doi.org/10.1038/s41586-020-2959-4

14. Ruis, P., Van Ly, D., Borel, V., Kafer, G. R., McCarthy, A., Howell, S., Blassberg, R., Snijders, A. P., Briscoe, J., Niakan, K. K., Marzec, P., Cesare, A. J., & Boulton, S. J. (2021). TRF2-independent chromosome end protection during pluripotency. Nature, 589(7840), 103–109. https://doi.org/10.1038/s41586-020-2960-y

15. Vessoni, A. T., Zhang, T., Quinet, A., Jeong, H.-C., Munroe, M., Wood, M., Tedone, E., Vindigni, A., Shay, J. W., Greenberg, R. A., & Batista, L. F. Z. (2021). Telomere erosion in human pluripotent stem cells leads to ATR-mediated mitotic catastrophe. The Journal of Cell Biology, 220(6). https://doi.org/10.1083/jcb.202011014

16. Herbig, U., Jobling, W. A., Chen, B. P. C., Chen, D. J., & Sedivy, J. M. (2004). Telomere Shortening Triggers Senescence of Human Cells through a Pathway Involving ATM, p53, and p21CIP1, but Not p16INK4a. Molecular Cell, 14(4), 501–513. https://doi.org/10.1016/S1097-2765(04)00256-4

17. Doksani, Y., & de Lange, T. (2016). Telomere-Internal Double-Strand Breaks Are Repaired by Homologous Recombination and PARP1/Lig3-Dependent End-Joining. CellReports, 17(6), 1646–1656. https://linkinghub.elsevier.com/retrieve/pii/S2211124716313948

18. Sim, X., Cardenas-Diaz, F. L., French, D. L., & Gadue, P. (2016). A Doxycycline-Inducible System for Genetic Correction of iPSC Disease Models. Methods in Molecular Biology, 1353(Chapter 179), 13– 23. http://link.springer.com/10.1007/7651_2014_179

19. Kimura, M., Stone, R. C., Hunt, S. C., Skurnick, J., Lu, X., Cao, X., Harley, C. B., & Aviv, A. (2010). Measurement of telomere length by the Southern blot analysis of terminal restriction fragment lengths. Nature Protocols, 5(9), 1596–1607. https://doi.org/10.1038/nprot.2010.124

20. Herbert, B.-S., Hochreiter, A. E., Wright, W. E., & Shay, J. W. (2006). Nonradioactive detection of telomerase activity using the telomeric repeat amplification protocol. In Nature Protocols (Vol. 1, Issue 3, pp. 1583–1590). https://doi.org/10.1038/nprot.2006.239

21. Henson, J. D., Lau, L. M., Koch, S., Martin La Rotta, N., Dagg, R. A., & Reddel, R. R. (2017). The C-circle assay for alternative-lengthening-of-telomeres activity. Methods (San Diego, Calif.), 114, 74–84. https://doi.org/10.1016/j.ymeth.2016.08.016

22. Cho, N. W., Dilley, R. L., Lampson, M. A., & Greenberg, R. A. (2014). Interchromosomal homology searches drive directional ALT telomere movement and synapsis. Cell, 159(1), 108–121. https://linkinghub.elsevier.com/retrieve/pii/S0092867414011003

23. Dilley, R. L., Verma, P., Cho, N. W., Winters, H. D., Wondisford, A. R., & Greenberg, R. A. (2016). Break-induced telomere synthesis underlies alternative telomere maintenance. Nature, 539(7627), 54–58. http://www.nature.com/articles/nature20099

24. Anderson, R., Lagnado, A., Maggiorani, D., Walaszczyk, A., Dookun, E., Chapman, J., Birch, J., Salmonowicz, H., Ogrodnik, M., Jurk, D., Proctor, C., Correia-Melo, C., Victorelli, S., Fielder, E., Berlinguer-Palmini, R., Owens, A., Greaves, L., Kolsky, K. L., Parini, A., … Passos, J. F. (2018). Length-independent telomere damage drives cardiomyocyte senescence. In bioRxiv (p. 394809). https://doi.org/10.1101/394809

25. Yu, Q., Katlinskaya, Y. V., Carbone, C. J., Zhao, B., Katlinski, K. V., Zheng, H., Guha, M., Li, N., Chen, Q., Yang, T., Lengner, C. J., Greenberg, R. A., Johnson, F. B., & Fuchs, S. Y. (2015). DNA-damage-induced type I interferon promotes senescence and inhibits stem cell function. Cell Reports, 11(5), 785–797. https://doi.org/10.1016/j.celrep.2015.03.069

26. Hewitt, G., Jurk, D., Marques, F. D. M., Correia-Melo, C., Hardy, T., Gackowska, A., Anderson, R., Taschuk, M., Mann, J., & Passos, J. F. (2012). Telomeres are favoured targets of a persistent DNA damage response in ageing and stress-induced senescence. Nature Communications, 3, 708. https://doi.org/10.1038/ncomms1708

27. Fumagalli, M., Rossiello, F., Clerici, M., Barozzi, S., Cittaro, D., Kaplunov, J. M., Bucci, G., Dobreva, M., Matti, V., Beausejour, C. M., Herbig, U., Longhese, M. P., & d’Adda di Fagagna, F. (2012). Telomeric DNA damage is irreparable and causes persistent DNA-damage-response activation. Nature Cell Biology, 14(4), 355–365. https://doi.org/10.1038/ncb2466

28. Sun, L., Tan, R., Xu, J., LaFace, J., Gao, Y., Xiao, Y., Attar, M., Neumann, C., Li, G.-M., Su, B., Liu, Y., Nakajima, S., Levine, A. S., & Lan, L. (2015). Targeted DNA damage at individual telomeres disrupts their integrity and triggers cell death. Nucleic Acids Research, 43(13), 6334–6347. https://doi.org/10.1093/nar/gkv598

29. Chen, J. (2016). The cell-cycle arrest and apoptotic functions of p53 in tumor initiation and progression. Cold Spring Harbor Perspectives in Medicine, 6(3), a026104. https://doi.org/10.1101/cshperspect.a026104

30. Vuoristo, S., Bhagat, S., Hydén-Granskog, C., Yoshihara, M., Gawriyski, L., Jouhilahti, E.-M., Ranga, V., Tamirat, M., Huhtala, M., Kirjanov, I., Nykänen, S., Krjutškov, K., Damdimopoulos, A., Weltner, J., Hashimoto, K., Recher, G., Ezer, S., Paluoja, P., Paloviita, P., … Kere, J. (2022). DUX4 is a multifunctional factor priming human embryonic genome activation. IScience, 25(4), 104137. https://doi.org/10.1016/j.isci.2022.104137

31. Hendrickson, P. G., Doráis, J. A., Grow, E. J., Whiddon, J. L., Lim, J.-W., Wike, C. L., Weaver, B. D., Pflueger, C., Emery, B. R., Wilcox, A. L., Nix, D. A., Peterson, C. M., Tapscott, S. J., Carrell, D. T., & Cairns, B. R. (2017). Conserved roles of mouse DUX and human DUX4 in activating cleavage-stage genes and MERVL/HERVL retrotransposons. Nature Genetics, 49(6), 925–934. https://doi.org/10.1038/ng.3844

32. Huang, Y., Liang, P., Liu, D., Huang, J., & Songyang, Z. (2014). Telomere regulation in pluripotent stem cells. Protein & Cell, 5(3), 194–202. https://doi.org/10.1007/s13238-014-0028-1

33. Sishc, B. J., Nelson, C. B., McKenna, M. J., Battaglia, C. L. R., Herndon, A., Idate, R., Liber, H. L., & Bailey, S. M. (2015). Telomeres and Telomerase in the Radiation Response: Implications for Instability, Reprograming, and Carcinogenesis. In Frontiers in Oncology (Vol. 5). https://doi.org/10.3389/fonc.2015.00257

34. Sato, N., Maehara, N., Mizumoto, K., Nagai, E., Yasoshima, T., Hirata, K., & Tanaka, M. (2001). Telomerase activity of cultured human pancreatic carcinoma cell lines correlates with their potential for migration and invasion. Cancer, 91(3), 496–504. https://doi.org/10.1002/1097-0142(20010201)91:3<496::aid-cncr1028>3.0.co;2-0

35. Varela, E., Schneider, R. P., Ortega, S., & Blasco, M. A. (2011). Different telomere-length dynamics at the inner cell mass versus established embryonic stem (ES) cells. Proceedings of the National Academy of Sciences of the United States of America, 108(37), 15207–15212. https://doi.org/10.1073/pnas.1105414108

36. Zalzman, M., Falco, G., Sharova, L. V., Nishiyama, A., Thomas, M., Lee, S.-L., Stagg, C. A., Hoang, H. G., Yang, H.-T., Indig, F. E., Wersto, R. P., & Ko, M. S. H. (2010). Zscan4 regulates telomere elongation and genomic stability in ES cells. Nature, 464(7290), 858–863. https://doi.org/10.1038/nature08882

37. Liu, L., Bailey, S. M., Okuka, M., Muñoz, P., Li, C., Zhou, L., Wu, C., Czerwiec, E., Sandler, L., Seyfang, A., Blasco, M. A., & Keefe, D. L. (2007). Telomere lengthening early in development. Nature Cell Biology, 9(12), 1436–1441. https://doi.org/10.1038/ncb1664

38. Bailey, S. M., Goodwin, E. H., Meyne, J., & Cornforth, M. N. (1996). CO-FISH reveals inversions associated with isochromosome formation. Mutagenesis, 11(2), 139–144. https://doi.org/10.1093/mutage/11.2.139

39. Bailey, S. M., Goodwin, E. H., & Cornforth, M. N. (2004). Strand-specific fluorescence in situ hybridization: the CO-FISH family. Cytogenetic and Genome Research, 107(1–2), 14–17. https://doi.org/10.1159/000079565

40. Yeager, T. R., Neumann, A. A., Englezou, A., Huschtscha, L. I., Noble, J. R., & Reddel, R. R. (1999). Telomerase-negative immortalized human cells contain a novel type of promyelocytic leukemia (PML) body. Cancer Research, 59(17), 4175–4179. https://www.ncbi.nlm.nih.gov/pubmed/10485449

41. Grobelny, J. V., Godwin, A. K., & Broccoli, D. (2000). ALT-associated PML bodies are present in viable cells and are enriched in cells in the G(2)/M phase of the cell cycle. Journal of Cell Science, 113 Pt 24, 4577–4585. https://doi.org/10.1242/jcs.113.24.4577

42. Henson, J. D., Cao, Y., Huschtscha, L. I., Chang, A. C., Au, A. Y. M., Pickett, H. A., & Reddel, R. R. (2009). DNA C-circles are specific and quantifiable markers of alternative-lengthening-of-telomeres activity. Nature Biotechnology, 27(12), 1181–1185. https://doi.org/10.1038/nbt.1587

43. Shlomovitz, I., Speir, M., & Gerlic, M. (2019). Flipping the dogma - phosphatidylserine in non-apoptotic cell death. Cell Communication and Signaling: CCS, 17(1), 139. https://doi.org/10.1186/s12964-019-0437-0

44. Liang, W., Qi, W., Geng, Y., Wang, L., Zhao, J., Zhu, K., Wu, G., Zhang, Z., Pan, H., Qian, L., & Yuan, J. (2021). Necroptosis activates UPR sensors without disrupting their binding with GRP78. Proceedings of the National Academy of Sciences of the United States of America, 118(39). https://doi.org/10.1073/pnas.2110476118

45. Saveljeva, S., Mc Laughlin, S. L., Vandenabeele, P., Samali, A., & Bertrand, M. J. M. (2015). Endoplasmic reticulum stress induces ligand-independent TNFR1-mediated necroptosis in L929 cells. Cell Death & Disease, 6(1), e1587. https://doi.org/10.1038/cddis.2014.548

46. McGrath, E. P., Centonze, F. G., Chevet, E., Avril, T., & Lafont, E. (2021). Death sentence: The tale of a fallen endoplasmic reticulum. Biochimica et Biophysica Acta (BBA) - Molecular Cell Research, 1868(6), 119001. https://doi.org/10.1016/j.bbamcr.2021.119001

47. Fernandez, R. J., Gardner, Z. J. G., Slovik, K. J., Liberti, D. C., Estep, K. N., Yang, W., Chen, Q., Santini, G. T., Perez, J. V., Root, S., Bhatia, R., Tobias, J. W., Babu, A., Morley, M. P., Frank, D. B., Morrisey, E. E., Lengner, C. J., & Johnson, F. B. (2022). GSK3 inhibition rescues growth and telomere dysfunction in dyskeratosis congenita iPSC-derived type II alveolar epithelial cells. ELife, 11. https://doi.org/10.7554/eLife.64430

48. Hosoi, T., Nakatsu, K., Shimamoto, A., Tahara, H., & Ozawa, K. (2016). Inhibition of telomerase causes vulnerability to endoplasmic reticulum stress-induced neuronal cell death. Neuroscience Letters, 629, 241–244. https://doi.org/10.1016/j.neulet.2016.07.027

49. Blasco, M. A., Lee, H. W., Hande, M. P., Samper, E., Lansdorp, P. M., DePinho, R. A., & Greider, C. W. (1997). Telomere shortening and tumor formation by mouse cells lacking telomerase RNA. Cell, 91(1), 25–34. https://doi.org/10.1016/s0092-8674(01)80006-4

50. Strong, M. A., Vidal-Cardenas, S. L., Karim, B., Yu, H., Guo, N., & Greider, C. W. (2011). Phenotypes in mTERT+/− and mTERT−/− mice are due to short telomeres, not telomere-independent functions of telomerase reverse transcriptase. Molecular and Cellular Biology, 31(12), 2369–2379. https://doi.org/10.1128/MCB.05312-11

51. Savage, S. A., & Bertuch, A. A. (2010). The genetics and clinical manifestations of telomere biology disorders. Genetics in Medicine: Official Journal of the American College of Medical Genetics, 12(12), 753–764. https://doi.org/10.1097/GIM.0b013e3181f415b5

52. Tummala, H., Walne, A., & Dokal, I. (2022). The biology and management of dyskeratosis congenita and related disorders of telomeres. Expert Review of Hematology, 15(8), 685–696. https://doi.org/10.1080/17474086.2022.2108784

53. Martínez, P., & Blasco, M. A. (2011). Telomeric and extra-telomeric roles for telomerase and the telomere-binding proteins. Nature Reviews. Cancer, 11(3), 161–176. https://doi.org/10.1038/nrc3025

54. Thompson, C. A. H., & Wong, J. M. Y. (2020). Non-canonical Functions of Telomerase Reverse Transcriptase: Emerging Roles and Biological Relevance. Current Topics in Medicinal Chemistry, 20(6), 498–507. https://doi.org/10.2174/1568026620666200131125110

55. Suram, A., & Herbig, U. (2014). The replicometer is broken: telomeres activate cellular senescence in response to genotoxic stresses. Aging Cell, 13(5), 780–786. https://doi.org/10.1111/acel.12246

56. Dan, J., Rousseau, P., Hardikar, S., Veland, N., Wong, J., Autexier, C., & Chen, T. (2017). Zscan4 Inhibits Maintenance DNA Methylation to Facilitate Telomere Elongation in Mouse Embryonic Stem Cells. Cell Reports, 20(8), 1936–1949. https://doi.org/10.1016/j.celrep.2017.07.070

57. Lee, K., & Gollahon, L. S. (2014). Zscan4 interacts directly with human Rap1 in cancer cells regardless of telomerase status. Cancer Biology & Therapy, 15(8), 1094–1105. https://doi.org/10.4161/cbt.29220

58. Dan, J., Zhou, Z., Wang, F., Wang, H., Guo, R., Keefe, D. L., & Liu, L. (2022). Zscan4 Contributes to Telomere Maintenance in Telomerase-Deficient Late Generation Mouse ESCs and Human ALT Cancer Cells. In Cells (Vol. 11, Issue 3, p. 456). https://doi.org/10.3390/cells11030456

59. Wang, F., Yin, Y., Ye, X., Liu, K., Zhu, H., Wang, L., Chiourea, M., Okuka, M., Ji, G., Dan, J., Zuo, B., Li, M., Zhang, Q., Liu, N., Chen, L., Pan, X., Gagos, S., Keefe, D. L., & Liu, L. (2012). Molecular insights into the heterogeneity of telomere reprogramming in induced pluripotent stem cells. Cell Research, 22(4), 757–768. https://doi.org/10.1038/cr.2011.201

