## Supplemental Information for "Telomeric DNA breaks in human induced pluripotent stem cells trigger ATR-mediated arrest and telomerase-independent telomere length maintenance"

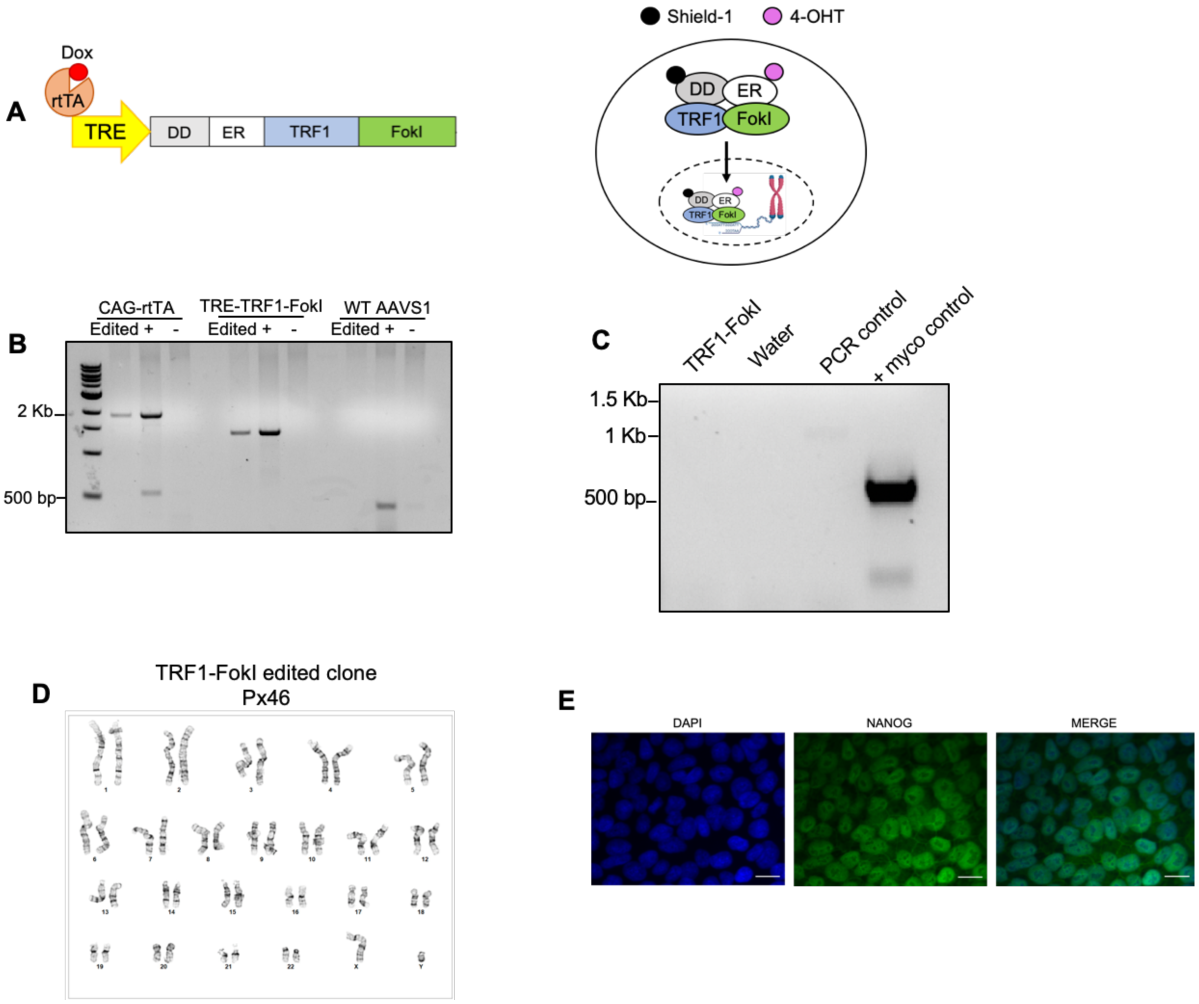


**Figure S1: Validation of engineered TRF1-FokI iPSC line**

- **A)** TRF1-FokI construct design. Upon addition of doxycycline, the constitutively expressed rtTA binds dox and drives transcription of the TRE-regulated DD-ER-TRF1-FokI allele. Once the protein is translated, Shield-1 ligand binds the DD domain to stabilize the cytosolic pool of protein and 4-OHT binds the ER domain to allow the stabilized protein to enter into the nucleus, where the TRF1 domain localizes the protein to telomere repeats to introduce telomeric double strand breaks.
- **B)** Genotyping PCR confirming the integration of CAG-rtTA and TRE-DD-ER-TRF1-FokI repair templates into the AAVS1 locus.
- **C)** PCR-based test to confirm the edited TRF1-FokI clone is negative for mycoplasma.
- **D)** The edited TRF1-FokI iPSC clone shows normal karyotype following the integration of rtTA and TRF1-FokI constructs into AAVS1.
- **E)** The edited TRF1-FokI iPSC clone maintains expression of Nanog, a marker of pluripotency. (Scale bar=10 $\mu$m).

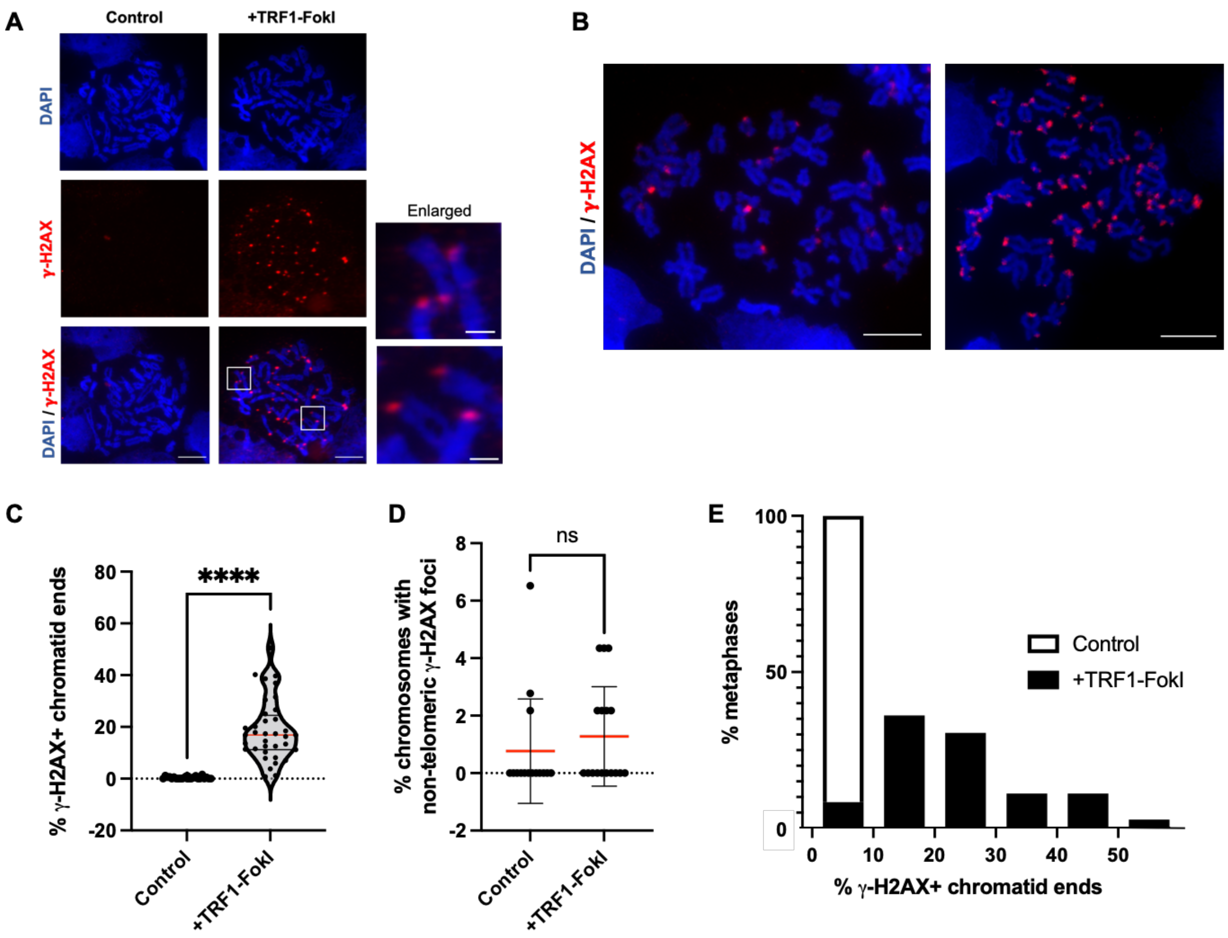


**Figure S2: TRF1-FokI DSBs are restricted to telomeres**

**A)** Immunofluorescent staining for **γ**-H2AX in uninduced cells and cells induced to express nuclear TRF1-FokI for 4 hours. Experimental setup is identical to that shown in Figure 1G. Left two panels: scale bar = 10 μM. Enlarged panels: scale bar = 2 μM.

**B)** Representative examples of metaphase spreads with varying frequencies of telomeric **γ**-H2AX foci. Scale bar = 10 μM.

**C)** Quantification of meta-TIF staining from 3 independent experiments. At least 30 spreads were analyzed per condition for a total of >1000 chromosomes. P-value is from two tailed unpaired t-test ($p<0.0001$).

**D)** Quantification of non-telomeric **γ**-H2AX foci in metaphase chromosomes stained as in C. Data represent the mean and SD from 3 independent experiments. P-value is from two tailed unpaired t-test ($p=0.419$).

**E)** Frequency distribution showing the percentage of metaphases analyzed that exhibited the indicated bin percentage of **γ**-H2AX positive chromatid ends (0-10%, 10-20% etc.). For example, 100% of control metaphases exhibited <10% **γ**-H2AX positive chromatid ends.

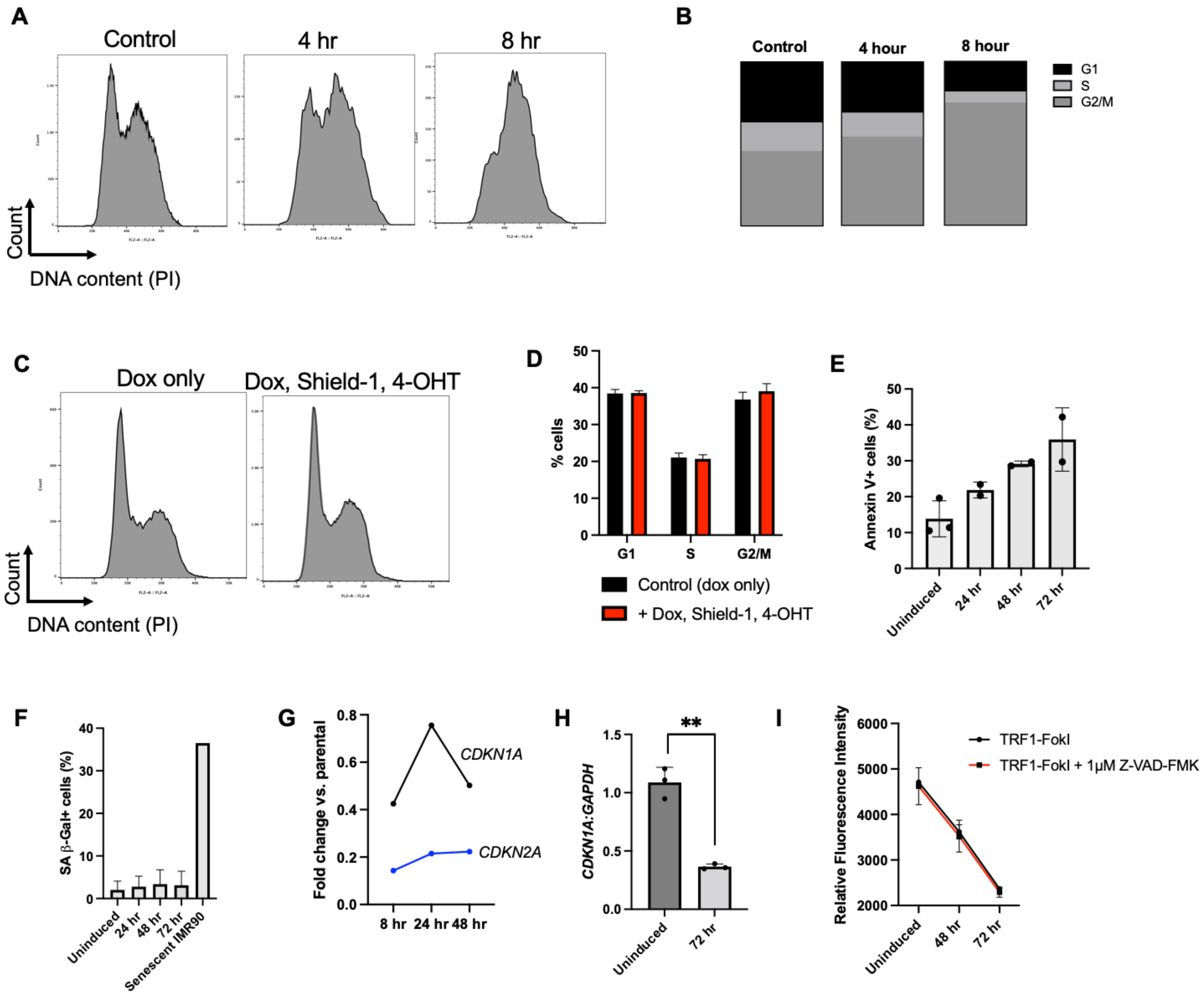


**Figure S3: Analysis of the effects of short- and long-term induction of TRF1-FokI**

- **A)** Representative histogram of DNA content of TRF1-FokI iPSCs uninduced (dox only) and induced (dox, Shield-1, 4-OHT) for 4 and 8 hours as measured by propidium iodide flow cytometry.
- **B)** Quantification of flow cytometry data from A. Data are representative of averages from 3 independent experiments.
- **C)** Representative histogram of DNA content of unedited parental SV20 iPSCs treated with dox only or complete induction medium containing dox, Shield-1, and 4-OHT as measured by propidium iodide flow cytometry.
- **D)** Quantification of flow cytometry data from C. Data are representative of average and SD from 2 independent experiments.
- **E)** Annexin V+ cells measured by flow cytometry across 48 hours of induction. Data represent the mean and SD from 2 independent experiments.

**F)** SA B-gal+ cells measured by flow cytometry across 72 hours of induction. IMR90 fibroblasts passaged to replicative senescence (px78) serve as a positive control (N=1). Data represent the mean and SD from 2 independent experiments.

**G)** Relative expression levels of senescence regulators p21 (CDKN1A) and p16 (*CDKN2A*) across 48-hour induction time course as measured by RNA seq compared to unedited parental cells.

**H)** Quantification of expression of *CDKN1A* mRNA transcripts at 72 hours of induction as measured by RT-qPCR and normalized to *GAPDH*. Data represent the mean and SD calculated from 3 independent experiments from 3 biological replicate samples, each containing triplicate qPCR reactions per time point.

**I)** Fluorometric cell viability assay of TRF1-FokI cells across 72 hours of induction with additional treatment with 1 μM pan-caspase inhibitor Z-VAD-FMK at the time of induction. Data represent the mean and SD from triplicate samples per condition measured at each time point.

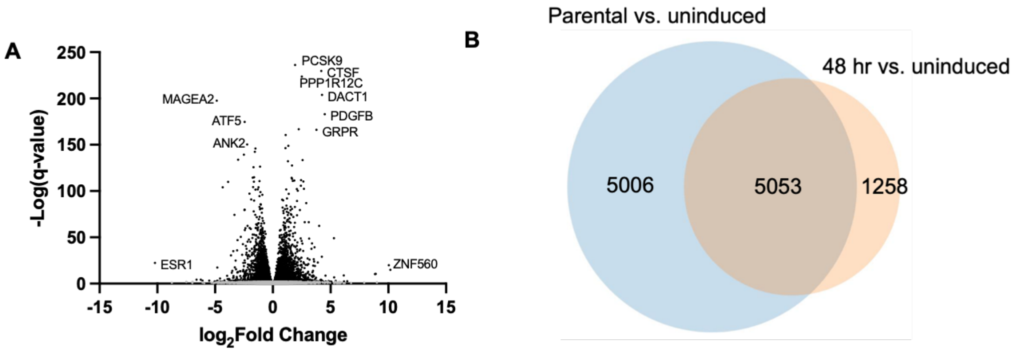


C

| Genes upregulated | | | Genes downregulated | | |
|---|---|---|---|---|---|
| Gene_name | log2FoldChange | padj | Gene_name | log2FoldChange | padj |
| PCSK9 | 1.926104305 | 7.111E-237 | MAGEA2 | -4.860260016 | 2.843E-198 |
| CTSF | 4.195884043 | 1.941E-230 | ATF5 | -2.441814385 | 1.81E-175 |
| PPP1R12C | 2.512843901 | 1.792E-224 | ANK2 | -2.225923793 | 3.942E-151 |
| DACT1 | 4.263586369 | 1.163E-204 | HSPA1A | -1.513927135 | 9.44E-147 |
| PDGFB | 4.493405431 | 8.217E-184 | SFPQ | -1.553237308 | 3.996E-143 |
| CCN2 | 2.233518126 | 1.362E-167 | NCOA5 | -2.502198496 | 6.513E-140 |
| GRPR | 3.770144836 | 7.907E-167 | MAP1A | -3.014631265 | 1.386E-134 |
| JADE1 | 1.10804512 | 3.107E-161 | UBTF | -1.108810136 | 5.396E-127 |
| PDLIM1 | 1.34083888 | 1.505E-149 | ODC1 | -1.681283275 | 1.619E-126 |
| HK1 | 1.097413117 | 3.895E-147 | BEND3 | -1.621610617 | 1.106E-115 |
| STARD8 | 2.57572245 | 3.543E-134 | HSPA1B | -1.738046018 | 2.768E-115 |
| PYGL | 1.30192692 | 6.886E-133 | HEXIM1 | -1.15003973 | 2.102E-111 |
| CORO2A | 1.658402829 | 2.389E-128 | PSAT1 | -1.648286471 | 8.476E-111 |
| SH2D3A | 1.62829841 | 4.779E-115 | CUZD1 | -3.866791799 | 1.225E-110 |
| HHLA1 | 1.663497688 | 4.748E-114 | UHRF1 | -1.706701413 | 5.801E-105 |
| BTG2 | 2.436143063 | 1.79E-112 | CSAG3 | -4.349278805 | 8.889E-105 |
| RAB20 | 1.86047669 | 3.671E-112 | RBM12 | -1.241075799 | 1.06E-104 |
| KCNK6 | 1.730349592 | 1.234E-111 | MAT2A | -1.310817804 | 1.433E-102 |
| NEDD9 | 2.63447555 | 5.818E-111 | ZBTB12 | -1.510975848 | 3.366E-102 |
| SLC16A3 | 2.42979308 | 1.901E-106 | DYNLL1 | -1.066730804 | 1.724E-99 |

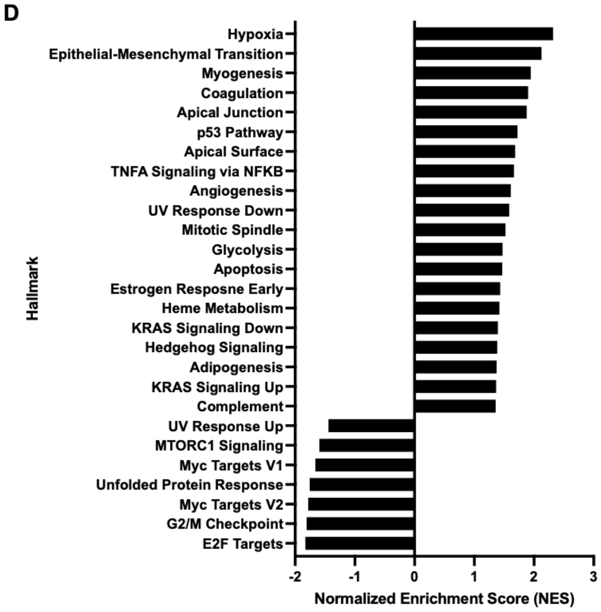


**Figure S4: Pathways differentially expressed in parental iPSCs cultured under induction conditions vs. uninduced TRF1-FokI iPSCs**

**A)** Volcano plot showing genes up- or down-regulated in unedited parental cells treated for 48 hours with dox, Shield-1, and 4-OHT compared to uninduced TRF1-FokI cells treated with dox only.

**B)** Venn diagram indicating the number of significantly (<0.05) differentially expressed lncRNA-, miRNA-, and protein-coding genes in each control condition. Based on these findings, drug-treated parental cells were selected as the control group for RNA-seq analyses.

**C)** Lists of the top 20 upregulated (left) and downregulated (right) genes in parental cells vs. uninduced TRF1-FokI iPSCs.

**D)** Gene set enrichment analysis (GSEA) of pathways significantly (<0.05) up- and down-regulated in parental vs. uninduced TRF1-FokI iPSCs.

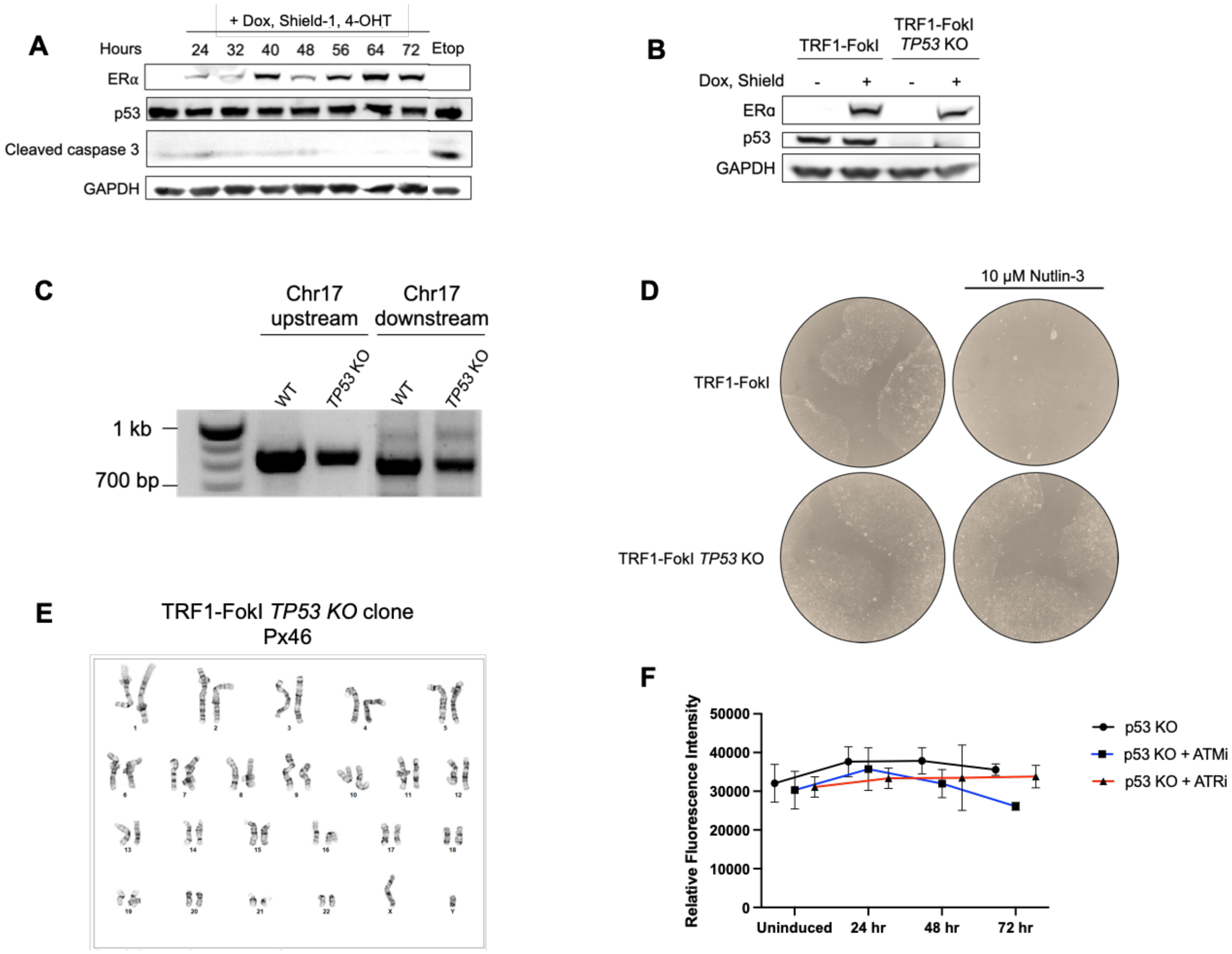


**Figure S5: Generation of TRF1-FokI *TP53* KO cells**

- **A)** Western blot analysis of total p53 protein levels in WT TRF1-FokI iPSCs across 96 hours of TRF1-FokI induction and in the same cells treated with 10 μM etoposide for 2 hours.
- **B)** Western blot confirmation of p53 knockout in TRF1-FokI iPSCs.
- **C)** Genotyping PCR of parental *TP53* WT TRF1-FokI line and *TP53 KO* clone using two primer sets designed against chromosome 17 upstream and downstream of the gRNA site in exon 10 of the *TP53* gene. Presence of a band in both PCRs confirms CRISPR deletion is contained within the TP53 coding sequence. For primer sequences see key resources table.
- **D)** Nutlin-3 viability assay to screen for loss of p53 function in expanded TRF1-FokI *TP53* KO cell line.
- **E)** Karyotype analysis of TRF1-FokI *TP53* KO edited clone. The cell line exhibits normal karyotype.
- **F)** Fluorometric cell viability assay of TRF1-FokI *TP53 KO* cells across 72 hours of induction with additional treatment with ATMi or ATRi. Data represent the mean and SD from triplicate samples per condition measured at each time point.

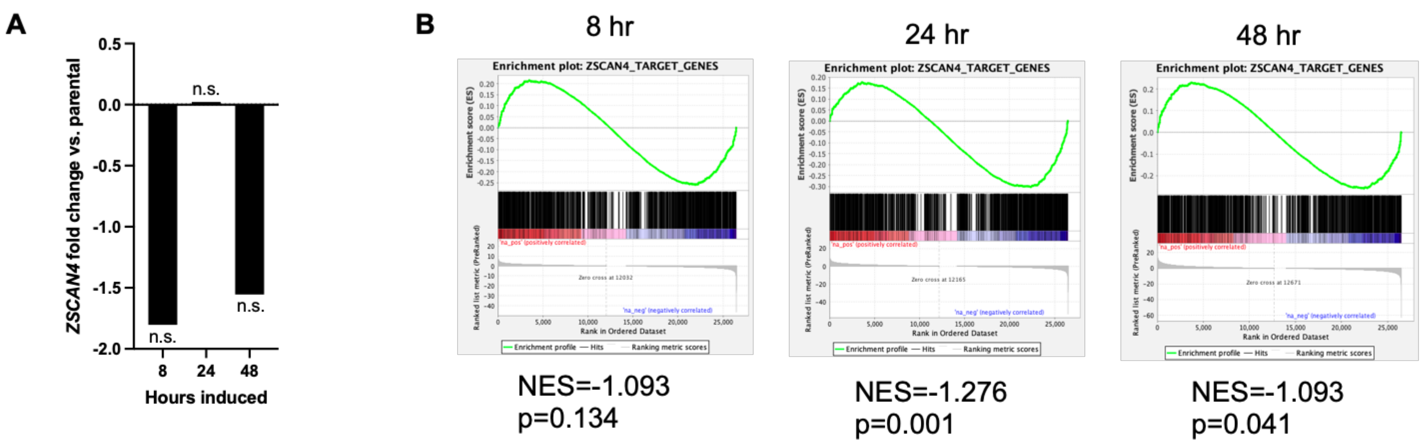


C

| Symbol (mouse) | Ortholog (human) | log2FoldChange (48 hr vs. parental) | p-value |
|---|---|---|---|
| Gm16429 | N/A | | |
| Inpp4b | INPP4B | -0.446883731 | 0.63401593 |
| Tdpoz4 | SPOP | 0.46467599 | 0.00183055 |
| Tcstv3 | N/A | | |
| Gm10696 | N/A | | |
| Zfp352 | KLF18 | -0.969487721 | 0.42094566 |
| Zscan4b | | | |
| Zscan4d | ZSCAN4 | -1.555680831 | 0.55764993 |
| Gm6351 | | | |
| Gm12794 | PRAMEF1 | Not detected | |
| AF067061 | N/A | | |
| Gm4303 | | | |
| Gm4302 | ALYREF | 0.414618569 | 0.00036532 |
| Spesp1 | SPESP1 | -1.326978957 | 0.77056715 |

**Figure S6: Human orthologues of mESC 2 cell-like genes are not enriched in TRF1-FokI induced iPSCs**

**A)** Relative *ZSCAN4* mRNA levels normalized to parental throughout 48 hours of TRF1-FokI induction.

**B)** GSEA indicating that the ZSCAN4 target gene set is significantly downregulated at 48 hours of TRF1-FokI induction.

**C)** Table indicating murine 2C-genes enriched in mESCs following deletion of TRF2 [13], with human orthologues and their relative expression as detected by RNA-seq at 48 hours of TRF1-FokI induction compared to parental unedited cells.

**A**

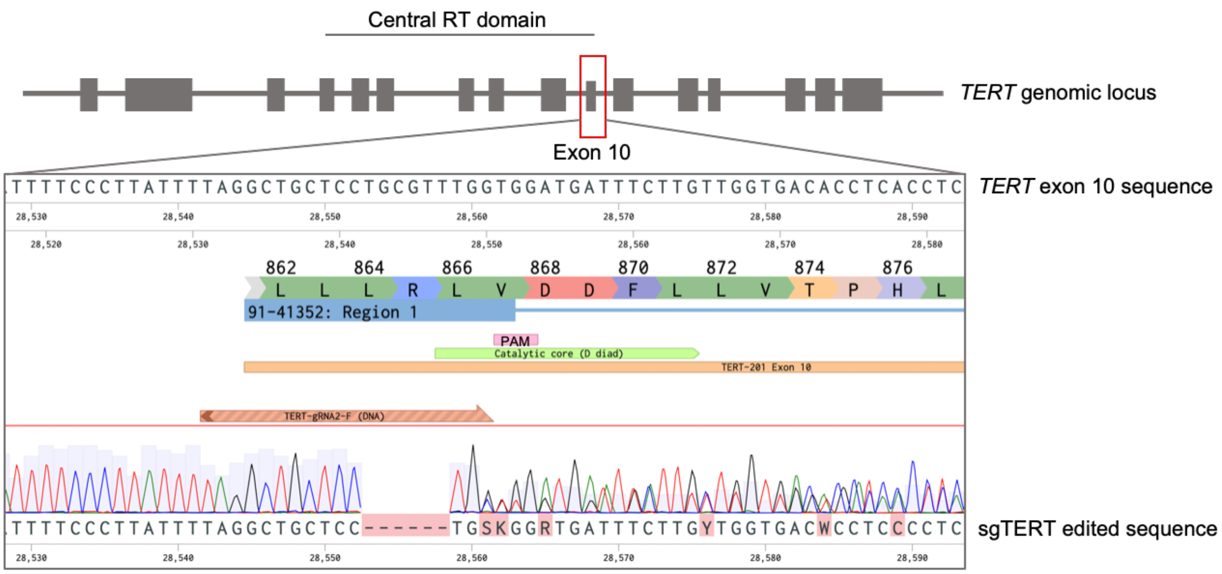


**B**

| | | | | | | | | | | | | | | | | | | | | | | | | |
|---|---|---|---|---|---|---|---|---|---|---|---|---|---|---|---|---|---|---|---|---|---|---|---|---|
| **Reference** | L | L | L | R | L | V | D | D | F | L | L | V | T | P | H | L | T | H | A | K | T | F | L | R | * |
| **Allele 1** | L | L | L | V | D | D | F | L | L | V | T | P | H | L | T | H | A | K | T | F | L | R | * | | |
| **Allele 2** | L | L | L | R | G | G | * | | | | | | | | | | | | | | | | | | |

**Figure S7: Generation of biallelic *TERT* mutant TRF1-FokI iPSCs**

**A)** CRISPR-based editing strategy for targeting exon 10 of the *TERT* locus in TRF1-FokI iPSCs and Sanger sequencing confirmation of an indel mutation adjacent to aspartate diads composing the catalytic core of the reverse transcriptase domain.

**B)** Amino acid sequence of resulting *TERT* alleles in CRISPR-targeted TRF1-FokI iPSCs.

**Table S1: Key resources and reagents used in this study**

| Reagent Type (Species) or Resource | Designation | Source or Reference | Identifiers | Additional Information |
|---|---|---|---|---|
| **Cell Lines** | | | | |
| iPS cell line (Homo sapiens) | PENN123i-SV20 | (Pashos et al. 2017) | | Parental |
| iPS cell line (Homo sapiens) | TRF1-FokI | This paper | | Engineered |
| iPS cell line (Homo sapiens) | TRF1-FokI *TP53* KO | This paper | | Engineered |
| iPS cell line (Homo sapiens) | TRF1-FokI sg*TERT* | This paper | | Engineered |
| Fibroblast cell line (Homo sapiens) | IMR-90 | (Nichols et al. 1977) | | |
| ALT- control cell line (Homo sapiens) | HEK-293T | (Graham et al. 1977) | | |
| ALT+ control cell line (Homo sapiens) | U2OS | (Ponten and Saksela 1964) | | |
| **Recombinant DNA Reagents** | | | | |
| Plasmids | pTRE-TIGHT-EGFP-donor | Paul Gadue | Addgene plasmid #22074 | |
| Plasmids | AAVS1-SA-2A-NEO-CAG-RTTA3 | Paul Gadue | Addgene plasmid #60431 | |
| Plasmids | PGK-AAVS1ZFNR | Paul Gadue | Addgene plasmid #60915 | |
| Plasmids | PGK-AAVS1ZFNL | Paul Gadue | Addgene plasmid #60916 | |
| Plasmids | pLenti-CMV-Puro-DEST-TRF1-FokI | Roger Greenberg | | |
| Plasmids | Px330 | | Addgene plasmid #42230 | |
| Plasmids | pCAG-SpCas9-GFP-U6-gRNA | | Addgene plasmid #79144 | |
| Primer | AAVS1-WT-F | (Sim et al. 2016) | | `CCC CTA TGT CCA CTT CAG GA` |
| Primer | AAVS1-WT-R | (Sim et al. 2016) | | `CAG CTC AGG TTC TGG GAG AG` |
| Primer | AAVS1-CAG-F | (Sim et al. 2016) | | `GAG CAT CTG ACT TCT GGC TAA TA` |
| Primer | AAVS1-CAG-R | (Sim et al. 2016) | | `GAA GGA TGC AGG ACG AGA AA` |
| Primer | AAVS1-TRE-F | (Sim et al. 2016) | | `GCA ATA GCA TCA CAA ATT TCA C` |
| Primer | AAVS1-TRE-R | (Sim et al. 2016) | | `GAA GGA TGC AGG ACG AGA AA (same as AAVS1-CAG-R)` |

| | | | | |
|---|---|---|---|---|
| Primer | Chr17 upstream F | This paper | | AGA GAC GAG GTT TCA TCA TGT T |
| Primer | Chr17 upstream R | This paper | | AAC AAC GTT CTG GTA AGG ACA A |
| Primer | Chr17 downstream F | This paper | | TTT TCA GTT GTG CCA GCT TCA T |
| Primer | Chr17 downstream R | This paper | | ACA GGA GAT GTT CTC ATA CAG GAG |
| Primer | TS | (Herbert et al. 2006) | | AAT CCG TCG AGC AGA GTT |
| Primer | ACX | (Herbert et al. 2006) | | GCG CGG CTT ACC CTT ACC CTT ACC CTA ACC |
| Primer | BranchedUniversal Primer | (Lai et al. 2016) | | [Phos] GAC TCT CAA CTA TC+T +A |
| Primer | G-Rich ONT | (Lai et al. 2016) | | [Phos] CCC TAA CCC TAA CCC TAA CCC TAA CCC TAA CCC TAA CCC TAG ATA GTT GAG AGT C |
| Primer | C-Rich ONT | This paper | | |
| gRNA | p53 gRNA | This paper | | AAT GAG GCC TTG GAA CTC A |
| gRNA | TERT gRNA | This paper | | TAG GCT GCT CCT CGT TTG G |
| **Chemicals and Small Molecules** | | | | |
| Chemical | Thiazovivin (TZV) | Cayman Chemical | CAT# 14245 | |
| Chemical | Puromycin (hydochloride) | Cayman Chemical | CAT# 13884 | |
| Chemical | G418 sulfate | Cayman Chemical | CAT# 13200 | |
| Chemical | Doxycycline (dox) | Cayman Chemical | CAT# 14422 | |
| Chemical | Shield-1 ligand | Aobious | CAT# AOB1848 | |
| Chemical | 4-OHT | Cayman Chemical | CAT# 17308 | |
| Chemical | Nutlin-3 | Cayman Chemical | CAT# 10004372 | |
| Chemical | Ku-55933 | Cayman Chemical | CAT# 16336 | |
| Chemical | VE-821 | Cayman Chemical | CAT #17587 | |
| Chemical | CCT241533 (hydrochloride) | Cayman Chemical | CAT# 19178 | |
| Chemical | LY2606368 | Cayman Chemical | CAT# 21490 | |
| Chemical | 5-Bromo-2'-deoxyuridine | Cayman Chemical | CAT# 15580 | |
| Chemical | 5-Bromo-2'-deoxycytidine | Thermo Fisher | CAT# AAJ6545603 | |
| Chemical | Hoechst | Thermo Fisher | CAT# 62249 | |
| Chemical | DAPI | Cayman Chemical | CAT# 14285 | |
| Chemical | Propidium iodide | Invitrogen | CAT# BMS500PI | |
| Chemical | Demecolcine solution | Sigma | CAT# D1925 | |
| Chemical | TriZOL | Invitrogen | CAT# 15596018 | |
| Chemical | Digoxigenin-11-dUTP | Sigma | CAT# 11093088910 | |

| Commercial Medias and Kits | | | | |
|---|---|---|---|---|
| Commercial Protein Product | Matrigel, Growth Factor Reduced | Corning | CAT# 354230 | Can use with or without phenol red |
| Commercial Media | StemMACS iPS-Brew XF | Miltenyi Biotec | CAT# 130-104-368 | |
| Commercial Media | mTeSR 1 | StemCell Technologies | CAT# 85850 | |
| Commercial Media | StemMACS passaging solution XF | Miltenyi Biotec | CAT# 130-104-688 | |
| Enzymatic Reagent | Accutase | Innovative Cell Technologies | CAT# AT104-500 | |
| Commercial Kit | CellEvent™ Senescence Green Flow Cytometry Assay Kit | Fisher Scientific | CAT# C10840 | |
| Commercial Kit | CellTiter-Blue® Cell Viability Assay | Promega | CAT# G8081 | |
| Commercial Kit | Lipofectamine™ Stem Transfection Reagent | Invitrogen | CAT# STEM00015 | |
| Commercial Kit | Gentra Puregene Cell Kit | Qiagen | CAT# 158745 | |
| Commercial Kit | Monarch RNA Cleanup Kit | NEB | CAT# T2040L | |
| Commercial Kit | Click-iT™ EdU Imaging Kit | Invitrogen | CAT# C10086 | |
| **Antibodies and In-Situ Probes** | | | | |
| Antibody | NANOG | Reprocell/Stemgent | CAT# 09-0020 | 1:250 4˚C overnight |
| Antibody | 53BP1 | Novus | CAT# NB100-304 | 1:250 4˚C overnight |
| Antibody | $\gamma$-H2AX | Millipore | CAT# 05-636 | 1:500 37˚C 1 hour |
| Antibody | Cleaved Caspase 3 (CC3) | Cell Signaling Technologies | CAT# 9664 | 1:1000 4˚C overnight |
| Antibody | Phospho-ATM (S1981) | R&D Systems | CAT# MAB22902-SP | 1:1000 4˚C overnight |
| Antibody | Total ATM | Cell Signaling Technologies | CAT# 2873T | 1:1000 4˚C overnight |
| Antibody | Phospho-ATR (Thr1989) | GeneTex | CAT# GTX128145 | 1:500 4˚C overnight |
| Antibody | Total ATR | ProteinTech | CAT# 19787-1-AP | 1:500 4˚C overnight |
| Antibody | Chk1 (FI-476) | Santa Cruz | CAT# sc-7898 | 1:1000 4˚C overnight |
| Antibody | Chk2 | Santa Cruz | CAT# sc-9064 | 1:1000 4˚C overnight |
| Antibody | TRF1 | Millipore | CAT# 04-638 | 1:500 4˚C overnight |
| Antibody | TRF1 | ProteinTech | CAT# 67592-1-Ig | 1:500 4˚C overnight |
| Antibody | Estrogen receptor alpha (ERa) | Abcam | CAT# ab16660 | 1:500 4˚C overnight |
| Antibody | GAPDH | Abcam | CAT# ab9485 | 1:15000 4˚C overnight |
| Antibody | Phospho-Histone H3 (AF488 conjugated) | Cell Signaling Tehcnologies | CAT# 3465S | 1:50 25˚C 90 min |
| Antibody | FITC Annexin V | BD Pharmingen | CAT# 560931 | 1:30 25˚C 30 min |
| Antibody | p53 | ProteinTech | CAT# 10442-1-AP | 1:1000 4˚C overnight |

| | | | | |
|---|---|---|---|---|
| Antibody | PML | Santa Cruz | CAT# sc-966 | 1:1000 4˚C overnight |
| PNA Probe | Cy3-Telo-C Probe | PNA Bio | CAT# F1002 | 0.5 µg/ml |
| PNA Probe | 488-Telo-G Probe | PNA Bio | CAT# F1008 | 0.5 µg/ml |
| Fab Fragments | Anti-Digoxigenin-AP, Fab fragments | Roche | CAT# 11093274910 | 1:20000 25˚C 30 min |
| Secondary Antibody | AlexaFluor 488 goat anti-rabbit IgG | Invitrogen | CAT# A-11034 | 1:500 25˚C 1 hour |
| Secondary Antibody | AlexaFluor 555 donkey anti-mouse IgG | Invitrogen | CAT# A-32773 | 1:500 25˚C 1 hour |
| Secondary Antibody | Rabbit anti-mouse HRP-conjugate | Abcam | CAT# ab97046 | 1:2000 25˚C 1 hour |
| Secondary Antibody | Goat anti-rabbit HRP-conjugate | Biorad | CAT# 170-6515 | 1:2000 25˚C 1 hour |
| **Molecular Biology Reagents** | | | | |
| Restriction Enzyme | CviAII | NEB | CAT# R0640L | |
| Restriction Enzyme | BfaI | NEB | CAT# R0568L | |
| Restriction Enzyme | MseI | NEB | CAT# R0525L | |
| Restriction Enzyme | NdeI | NEB | CAT# R0111L | |
| Recombinant Enzyme | rSAP | NEB | CAT# M0371L | |
| Recombinant Enzyme | T4 DNA ligase | NEB | CAT# M0202S | |
| Molecular Biology Reagent | CDP-Star | Roche | CAT# 11759051001 | |
| Modified Nucleotide | DIG-11-dUTP | Roche | CAT# 11558706910 | |
| Molecular Biology Reagent | DIG Easy Hybridization Granules | Roche | CAT# 11796895001 | |
| Molecular Biology Reagent | Blocking Reagent | Roche | CAT# 11096176001 | |
| Recombinant DNA Marker | DIG Labeld DNA Molecular Weight Marker II | Roche | CAT# 11218590910 | |
| Molecular Biology Reagent | Hybond-XL | Cytiva/Amersham | CAT# RPN303S | |
| Recombinant Enzyme | Klenow Fragment | NEB | CAT# M0212S | |
| Recombinant Enzyme | Lambda Exonuclease | NEB | CAT# M0262S | |
| Recombinant DNA Polymerase | Go-TAQ Flexi | Promega | CAT# M8298 | |
| Recombinant DNA Polymerase | Phi29 DNA polymerase | NEB | CAT# M0269 | |
| Recombinant Enzyme | Exonuclease I | NEB | CAT# M0293S | |

| | | | | |
|---|---|---|---|---|
| Recombinant Enzyme | Exonuclease III | Promega | CAT# M1815 | |
| Recombinant Enzyme | RNase A | Qiagen | CAT# 19101 | |
| Recombinant Enzyme | Terminal transferase | NEB | CAT# M0315S | |
| **Software** | | | | |
| Image Analysis Software | FIJI | NIH | RRID:SCR_002285 | https://imagej.net/Fiji |
| Software | Benchling | | | benchling.com |
| Software | GraphPad Prism 9.2 | | | |
| Software | Salmon | | | |
| Software | GSEA | | | |
| Flow Cytometry Analysis Software | FlowJo | BD Biosciences | RRID:SCR_008520 | |
| Image Analysis Software | ImageQuant TL 8.2 | Cytiva | RRID:SCR_018374 | |
